# Scalable expansion and hepatic zone maturation of hepatic progenitor cells from human pluripotent stem cells

**DOI:** 10.1101/2025.06.13.659636

**Authors:** Mina Ogawa, Jeff C Liu, Abolfazl Dadvar, Britney Tian, Xinyuan Zhao, Ian Fernandes, Kentaro Minegishi, Kenichiro Takase, Changyi Cui, Marcela Hernandez, Yuichiro Higuchi, Hiroshi Suemizu, Ian McGilvray, Sonya MacParland, Gary Bader, Shinichiro Ogawa

## Abstract

Scalable generation of functionally mature hepatocytes from human pluripotent stem cells (hPSCs) is vital for regenerative medicine, disease modeling, and drug screening. We present a modular differentiation platform that enables serial expansion and cryopreservation of hepatic progenitors (hepatoblasts; HBs), supporting efficient downstream maturation. By modulating bile acid signaling with FGF19 and culturing under hypoxia, we preserved HB proliferation and bipotency. Subsequent thyroid hormone treatment induced alpha-fetoprotein-negative, albumin-positive hepatocytes, while WNT pathway modulation promoted zonal identity, mimicking *in vivo* hepatic metabolic heterogeneity. Zonation was confirmed by differential drug-metabolizing enzyme activity, mitochondrial function, and enhanced engraftment in a liver injury mouse model. To evaluate translational relevance, mature hepatocytes were incorporated into extrusion-based 3D bioprinted constructs, which maintained hepatic viability and function. This integrated strategy of combining progenitor cell banking, zonation control, and tissue engineering offers a scalable and clinically relevant approach for generating functional human liver tissue suitable for therapeutic development.

## Introduction

Liver transplantation remains the only definitive treatment for patients with end-stage liver failure^1^. However, its clinical application is significantly constrained by a global shortage of donor organs and the lifelong immunosuppression required to maintain graft tolerance^2^. These limitations highlight an urgent need for alternative regenerative strategies, including cell-based and tissue-engineered therapies, that can restore or replace hepatic function^3–5^.

However, developing these alternatives poses considerable challenges, particularly due to the extensive cellular and functional demands of the liver. As the largest internal organ, the liver comprises over 100 billion cells and executes more than 500 essential physiological functions, including xenobiotic detoxification, plasma protein synthesis, glucose and lipid metabolism, and bile acid production^6^. Hepatocytes, the primary functional units of the liver, account for approximately 80% of the parenchymal cell population^7^. Assuming that vital hepatic functions could be sustained with as little as over 1% of the total adult liver cell mass, it is estimated that a minimum of 1 × 10⁹ functional hepatocytes would be required to confer therapeutic benefit. This estimate is based on the relatively consistent hepatocellular density observed across mammalian species (∼100–120 × 10⁶ cells per gram of liver tissue) and the average adult human liver mass of 1–1.5 kg^8,9^. In some cases, an even greater number of cells may be necessary to achive clinical efficiency. Yet, reliably generating such vast quantities in a scalable and cost-effective manner remains a major hurdle.

One promising avenue involves the use of human pluripotent stem cells (hPSCs), including embryonic stem cells and induced pluripotent stem cells, which offer a theoretically unlimited and renewable source of liver cells^10^. Importantly, recent advances in clinical trials have demonstrated significant progress in their therapeutic application^11–13^. Despite this promise, two major limitations persist in the context of liver therapy: (1) the inefficiency of generating fully mature and functional hepatocytes, and (2) the inability to scale cell production beyond the pluripotent stage without significant losses in yield, cellular identity, or lineage fidelity^14–16^. Most current differentiation protocols suffer a marked drop in yield after definitive endoderm induction. Additionally, many liver organoid models depend heavily on mitogenic cues (e.g., HGF, EGF) or undefined ECM (Extra cellular Matrix), both of which may drive off-target differentiation, reduced structural fidelity, and significantly increase production complexity^17–22^.

Although recent advances such as small molecules expansion, transcriptional reprogramming, and direct conversion have facilitated the derivation of hman liver cells, they still fall short of replicating the full functional repertoire of native hepatocytes^23,24^. Issues such as batch-to-batch variability, functional decline during maturation, and limited long-term stability continue to pose significant challenges. While some engineered liver tissues have shown efficacy in preclinical transplantation models, key obstacles remain, including the identification of specific hepatocyte clones suitable for large-scale culture, and the development of immunoevasive strategies to prevent allogeneic rejection^25,26^.

In this study, we report a chemically defined, scalable platform for the expansion and maturation of hepatic progenitor cells, also called hepatoblasts (HBs) derived from hPSCs. By modulating bile acid signaling pathways under hypoxic conditions within specialized media, we achieved continuous passaging and cryopreservation of ALB⁺/AFP⁺ bipotent progenitors with minimum lineage instability. These HBs retained their hepatic identity and were readily transitioned into 3D aggregates/organoid cultures in either suspension or ECM-supported systems to generate functional hepatocytes and ciliated cholangiocytes in fully defined media. Importantly, thyroid hormone supplementation promoted hepatic maturation, yielding ALB-only hepatocytes which are characteristic of more advanced developmental stages. Furthermore, zonal metabolic heterogeneity, a hallmark of native liver architecture, was successfully recapitulated through small-molecule modulation of the WNT signaling pathway. Together, these advances lay the groundwork for large-scale, clinically relevant production of mature, functionally zonated hPSC-derived liver cells, opening new avenues for regenerative therapy in liver disease.

## Results

### FGF19 signaling, along with the activation of Wnt pathway and inhibition of TGF-β promotes the proliferation of hPSC-derived hepatoblasts

To facilitate the large-scale manufacturing of hepatocytes derived from hPSCs, we first evaluated their proliferative potential within our existing differentiation protocol^27^. Following endoderm differentiation, we employed flow cytometry analysis to assess the presence of ALB positive (ALB^+^) cells. As shown in Figure 1a, the frequency of ALB^+^ cells increases dramatically over the 14-day culture period. On the other hand, Ki67^+^ cells exhibited a declining trend throughout the differentiation, as assessed by flow cytometry, with percentages of 19.85 % ± 1.52, 5.37% ± 0.46, and 1.73% ± 0.23 (mean ± SEM) on day 15, day 21/23, and day 29, respectively (Fig. 1a, b). Consistent with the flow cytometry analysis, the differentiated cells displayed significantly upregulation of hepatocyte-related genes, including *ALB*, *AFP*, *HNF4a*, *CYP7A1*and *CYP3A7* (Fig. 1c).

**Figure 1.**
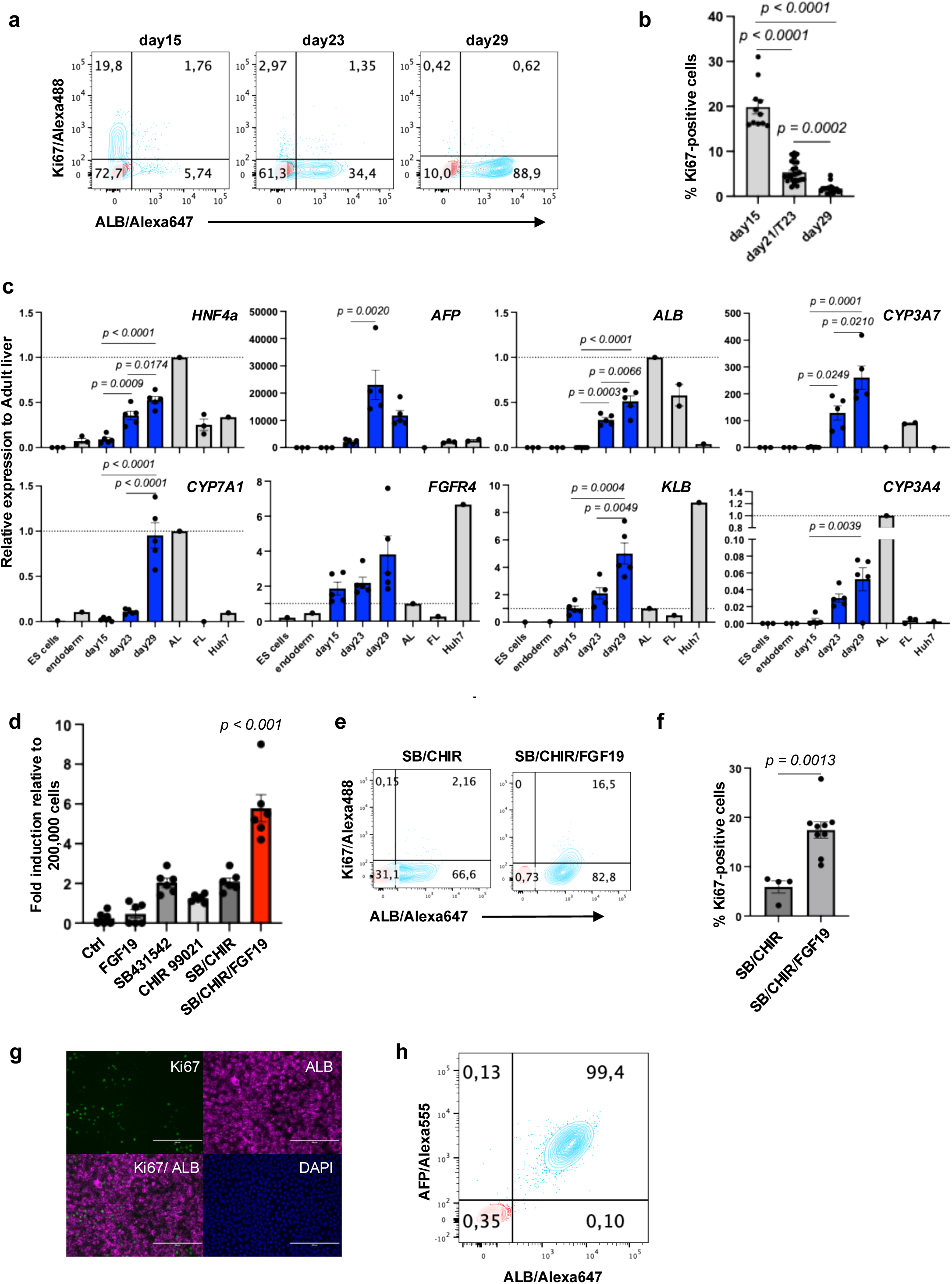
FGF19 signaling, along with Wnt activation and TGF β inhibition, promotes proliferation of hPSC-derived hepatoblasts. a) Flow cytometry analysis showing the proportion of Ki67+, and ALB+ cells at day 15, day23, and day29 of differentiation. b) Quantification of Ki67 positive cells in day 15, day 21or 23, and day 29 H9 hepatoblasts by flow cytometry. One-way ANOVA. DATA are represented as mean ± SEM (*n* = 11 for day 15, *n* = 25 for day 21 or 23, *n* = 19 for day 29). c) RT-qPCR analysis of the expression of the indicated gene in each population at different stages (ES cells, endoderm, day 15, day 23, day 29) of H9-derived hepatoblasts. AL adult liver, FL fetal liver. One-way ANOVA. DATA are represented as mean ± SEM (*n* = 5). d) Quantification of cells in day 29+6 H9 hepatoblasts after treatment with different indicated cytokine/ small molecule combinations. SB/CHIR/FGF19 group showed significantly higher values compared to all other groups (p< 0.001). One-way ANOVA. DATA are represented as mean ± SEM (*n* = 6). e) Flow cytometry analysis showing the proportion of Ki67+ and ALB+ cells at day 29+6 of differentiation in SB/CHIR or SB/CHIR/FGF19. f) Quantification of Ki67 positive cells in day 29+6 H9-derived hepatoblasts after treatment with SB/CHIR or SB/CHIR/FGF19. Two-tailed Student’s test (*n* = 4 and 9). g) Microscopic analysis showing the co-expression of ALB (pink) and Ki67 (green) in H9-derived hepatoblasts. Scale bar: 200µm. h) Flow cytometry analysis showing the proportion of ALB+ and AFP+ cells at day 29+6 of differentiation under SB/CHIR/FGF19 treatment.

CYP7A1 is a rate-limiting enzyme in the bile acid synthesis pathway. Bile acid signalling plays a pivotal role in mediating this function, in part through the effect of fibroblast growth factor 19 (FGF19), an enterokine secreted by the intestinal epithelium in response to bile acid flux^28,29^. The upregulation of CYP7A1 is critical for hepatocyte development, liver homeostasis, and regeneration as demonstrated by the finding that overexpression of the enzyme impairs regeneration following partial hepatectomy^30^. FGF19 exerts its effects on hepatocytes through interaction with the FGFR4/β-Klotho receptor complex both of which were found to be expressed in day 29 hPSC-derived hepatoblasts (Fig. 1c). The expression of this receptor complex in HBs suggested that FGF19/FGFR4/β-Klotho signaling may impact the growth of these cells^31^. To test this, we treated day 29 cells with FGF19 alone or in combination with the TGF-β pathway inhibitor SB431542 and the WNT agonist CHIR99021. Although FGF19 alone did not significantly stimulate proliferation, its combination with SB431542 and CHIR99021 led to a marked enhancement of HB proliferation, resulting in a six-fold increase in total cell number (Fig. 1d). Importantly, FGF19 signaling led to a concentration-dependent reduction of *CYP7A1* expression in day 29 HBs, accompanied by an upregulation of the proliferation-associated transcription factor *FOXM1* (Supplementary Fig. 1). FGF19 treated population contained a higher proportion of proliferating cells as demonstrated by Ki67 expression. These proliferating cells retain expression of ALB and AFP, indicating preservation of hepatic progenitor identity during expansion (Fig. 1e– h). Collectively, these results demonstrate that modulation of bile acid signaling via FGF19, in conjunction with TGF-β inhibition and WNT activation, effectively induces proliferation of hPSC-derived hepatoblasts while maintaining their bipotent hepatic progenitor phenotype.

### The hPSC-derived hepatic progenitor population can be serially expanded and cryopreserved

Given that the minimum estimated cell dose required for treating liver failure exceeds one billion functional hepatocytes, it is imperative to establish robust methods for large-scale cell production. Ideally, strategies that enable the expansion of mid- to late-stage differentiated hepatic populations are preferred, as they reduce the reliance on repeated expansion and differentiation of hPSCs for each production cycle. Consequently, we investigated whether HBs cultured with a combination of a TGFβ inhibitor SB 431542 (SB), a Wnt agonist CIHR 99021 (CHIR), and FGF19, hereafter referred to as SCF19, could be further expanded. Under SCF19 culture conditions, HBs seeded at a density of 100,000 cells per one well in a 12-well plate proliferated and increased in number by 6- to10-fold over a period of 8 to 10 days (Fig. 2a). Flow cytometry analysis showed that the expanded population co-expressed ALB and AFP, indicating that the cells retained their hepatic progenitor profile (Fig. 2b). The proliferation rate gradually declined with repeated passage under normal oxygenation conditions, and after the fourth passage, the increase in cell number was only 2-fold. However, when the cultures were maintained under hypoxic conditions, the proliferative potential of the cells was preserved, enabling up to ten consecutive passages. Under hypoxic conditions, it was possible to expand the HB population by 1000-fold over five consecutive passages (total of 53 days) (Figure 2a) generating 2000 hepatoblasts from a single hPSC. Flow cytometric analysis demonstrated that the proportion of ALB^+^/Ki67^+^ proliferating HBs decreased from 57.96% ± 1.78 (mean ± SEM) on day 2 post-passage 1 (P1) to 8.07% ± 0.85 by day 10. A similar pattern was observed in passage 2 (P2), with the ALB^+^/Ki67^+^ population declining from 55.05% ± 2.67 on day 2 to 3.06% ± 1.21 by day 10. By passage 3 (P3), a modest reduction in proliferative capacity was observed, with the ALB^+^/Ki67^+^ population measuring 23.98% ± 0.99 as early as day 2 post-passage (Supplementary Fig. 2a). Importantly, flow cytometry analysis further confirmed that the ALB^+^/AFP^+^ double-positive hepatic progenitor phenotype was consistently maintained throughout multiple passages (Fig. 2b). Immunohistochemistry analysis indicated that the ALB^+^/AFP^+^ hepatic progenitor cells were proliferative as indicated by Ki67 co-positivity (Figure 2c). Maintenance of this progenitor phenotype was dependent on culture in SCF19, as cells cultured in conventional hepatocyte differentiation conditions (e.g., DEX and OSM) lost expression of ALB and AFP (Supplementary Fig. 2b).

**Figure 2.**
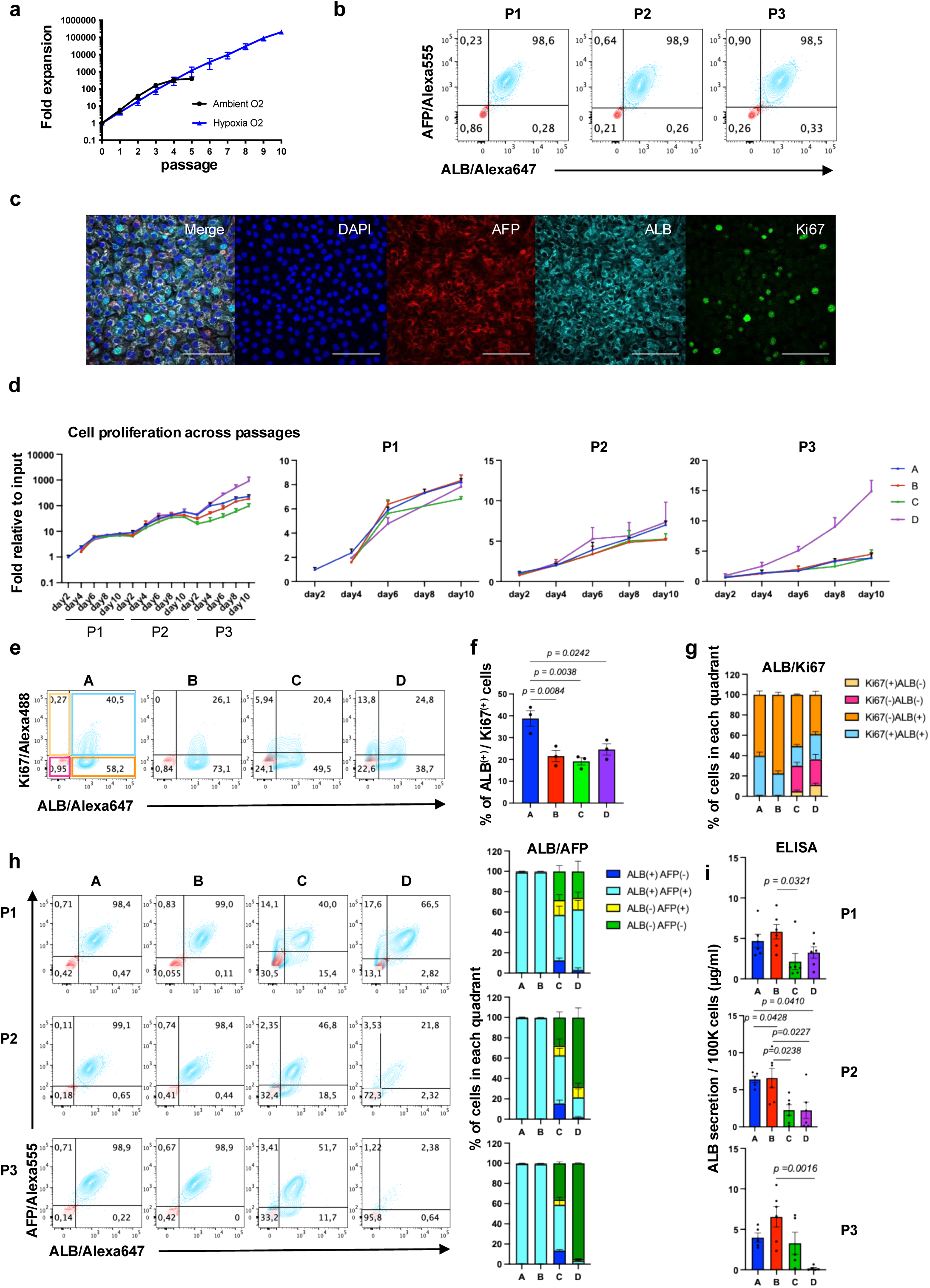
hPSC-derived hepatic progenitors can be serially expanded. **a)** Quantification of H9-derived hepatoblast numbers under hypoxic or ambient conditions over the passages. b) Flow cytometry analysis showing the proportion of ALB+ and AFP+ cells at each passage cultured in SCF19 expansion cocktails. P indicates the passage number during cell culture. c) Immunostaining analysis showing the co-expression of ALB (cyan), AFP (red), and Ki67 (green) in H9-derived hepatoblasts. Scale bar: 50µm. d) Quantification of fold change in H9-derived hepatoblast numbers cultured in different expansion cocktails. P indicates the passage number during cell culture. e) Flow cytometry analysis showing the proportion of Ki67+ and ALB+ cells at day 6 of passage 1 cultured in different expansion cocktails. f) Quantification of ALB+ and Ki67 + cells at day 6 of passage 1 cultured in different expansion cocktails. One-way ANOVA (*n* = 3). g) Quantification of positive cells in each quadrant, indicated by colours in Figure (e) (*n* = 4). h) Flow cytometry analysis and its quantification of ALB+ and AFP+ populations across different expansion cocktails over three passages. The left panel show representative flow cytometry profiles. The right panels display the corresponding quantification of ALB+ and AFP+ cell populations (*n* = 3). i) ELISA analysis showing ALB secretion in different cocktails, as indicated in Figure (h). One-way ANOVA (*n* = 5-6).

The SCF19-expanded HBs could be cryopreserved at all passages, with post-thaw cell viability consistently exceeding 85%. The cryopreserved HBs retained their proliferative capacity and hepatic identity through extended passaging post-thaw. Karyotype analysis confirmed genomic stability at later passage (P5) following recovery from cryopreservation (Supplementary Fig. 2c), further supporting the suitability of this system for long-term cell banking and downstream applications.

Recent studies have shown the feasibility of expanding and serially passaging hPSC-derived hepatoblasts using specific cytokine combinations^20,32^. We compared our designed media A (SB, CHIR, FGF19) and B (SB, CHIR, FGF19 + RI), with established cocktails, medium C (SB, CHIR, HGF, EGF)^20^ and medium D (SB, CHIR, bFGF, RI)^32^, to assess their capacity to propagate HBs. All four media formulations supported HB proliferation and population expansion through three serial passages (P3) (Fig. 2d). However, only the cultures maintained in media A (SCF19) and B (SCF19 +RI) consistently retained a high percentage of ALB^+^/AFP^+^ HBs across passages. Among these, expansion medium A exhibited the highest proportion of ALB⁺/Ki67⁺ cells on day 6 of P1 (Fig. 2e-g). Flow cytometry analysis of ALB and AFP on day 6 of P1 revealed the following percentages of ALB ^+^/ AFP^+^ cells: medium A (98.77 % ± 0.19), medium B (99.03 % ± 0.03), medium C (45.67 % ± 8.35), and medium D (59.87 % ± 8.98) (mean ± SEM). Notably, ALB^-^/ AFP^-^ cells were observed in medium C (27.20 % ± 5.37) and medium D (26.33 % ± 9.90), and these double negative populations expanded further in subsequent passages (Fig. 2h). These findings highlight the differential impact of various expansion cocktails on both proliferative capacity and phenotypic stability of hPSC-derived hepatoblasts. Immunohistochemical analysis further supported this observation: populations cultured under conditions A and B consisted predominantly of ALB^+^/ AFP^+^ cells, whereas those cultured under conditions C and D contained ALB^-^/ AFP^-^ cells (Supplementary Fig. 2d). These phenotypic differences were corroborated by functional assays that showed significantly higher levels of albumin secretion in cells cultured in conditions A and B than those cultured in conditions C and D during the first passage (Fig. 2i). Similar trends were observed in suspension culture, where aggregated cells maintained after initial monolayer expansion (P1) and subsequent passage (P2), in each respective medium exhibited higher albumin secretion under conditions A and B. (Supplementary Fig. 2e). Together, these findings show that SCF19 medium are effective in supporting the propagation of hPSC-derived ALB^+^/ AFP^+^ hepatic progenitor cells.

### Generation of organoids in SCF19 medium

Optimizing the expansion culture conditions for hepatic progenitor cells is paramount for their efficient formation of organoids. In this study, we explore the incredible potential of generating organoids using SCF19 medium. Notably, we showcase the ability to directly thaw cryopreserved HBs and seed them into microwell arrays to form 3D organoids. Using the microwell plate, HBs were seeded at varying densities ranging from 20 to 1000 cells per microwell (Fig. 3a, b). Over time, the 3D aggregates exhibited increased surface area and enhanced ALB secretion, with secretion plateauing around day 8 at higher seeding densities (500 and 1000 cells per microwell), consistent with confluence-induced growth arrest.

**Figure 3.**
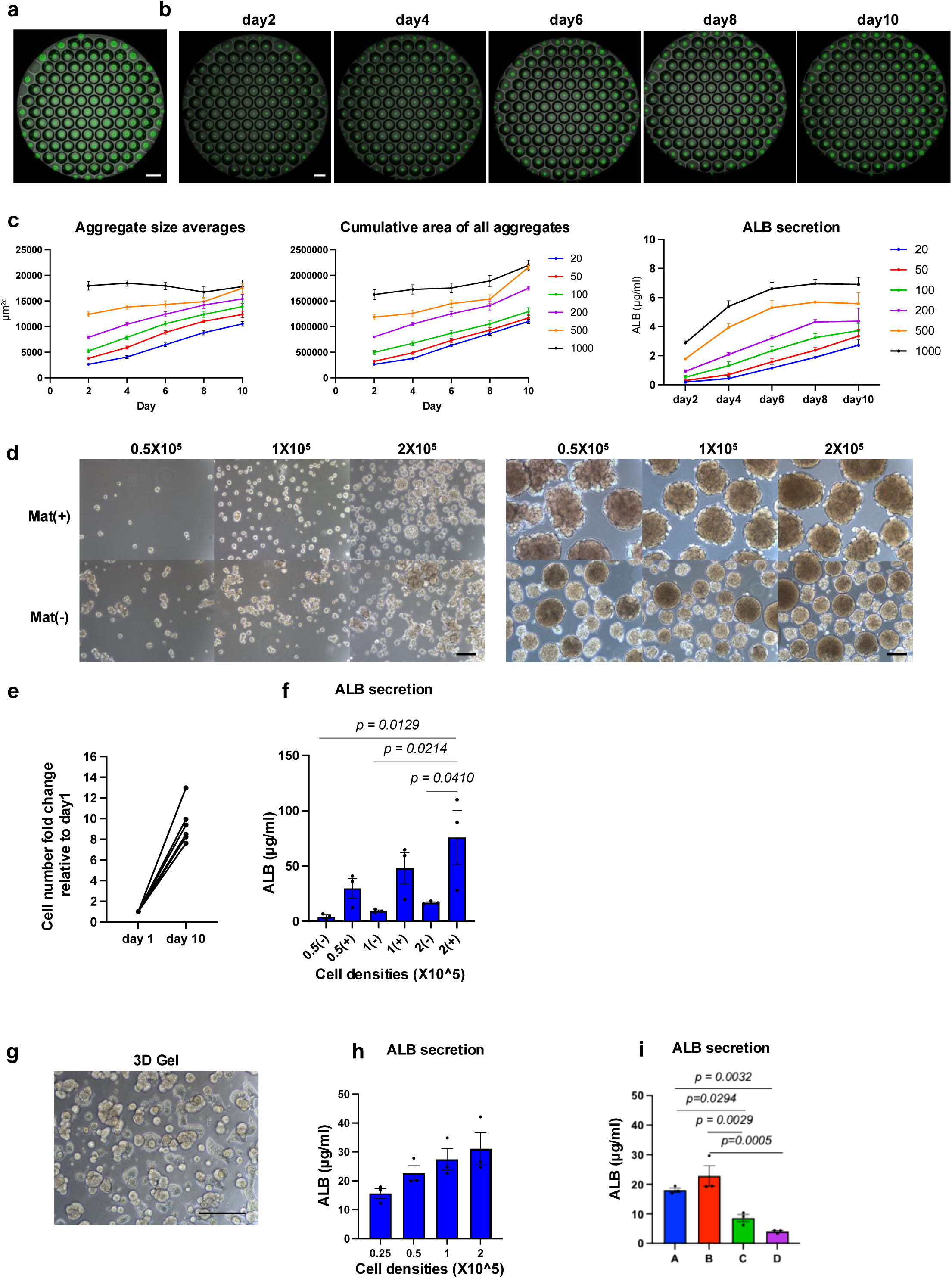
SCF19 Medium supports robust generation of 3D hepatic aggregates from hPSCs. a) Microscopic analysis of GFP-H9-derived 3D hepatoblasts organoids in microwells. Scale bar: 500µm. b) Microscopic analysis showing the growth in cryopreserved GFP-H9-derived 3D hepatoblasts organoids in microwells over a 10-day period. Scale bar: 500µm. c) Quantification of aggregate size averages (left), cumulative area of all aggregates (middle), and ALB secretion (right) of H9-derived hepatoblasts aggregate cultures with different initial cell numbers over a 10-day period. d) Microscopic analysis of H9-derived 3D hepatoblasts organoids in suspension culture with or without Matrigel. Images were taken on day 2 (left) and day 10 (right). Scale bar: 100µm. e) Quantification of cell numbers in 3D aggregates over 10 days. (*n* = 6). f) ALB secretion in H9-derived 3D hepatoblasts suspension culture at different cell densities, with or without Matrigel. One-way ANOVA (*n* = 3). g) Microscopic image of H9-derived 3D hepatoblasts organoids in Matrigel. Scale bar: 250µm. h) ALB secretion in H9-derived 3D hepatoblasts in Matrigel culture at different cell densities (*n* = 3). i) ALB secretion in 3D hepatoblasts cultured in Matrigel with different expansion cocktails. One-way ANOVA (*n* = 3).

Interestingly, lower-density microwells supported more robust cell proliferation, likely due to the availability of free space, as confirmed by ALB secretion profiles (Fig. 3b, c and Supplementary Fig. 3a). The ability to generate 3D aggregates from single cells was also confirmed through direct passaging (Supplementary Fig. 3b, c). Similarly, cryopreserved HBs resumed proliferation when thawed and cultured as single-cell suspensions at varying densities. The addition of 0.75% (v/v) Matrigel to the suspension culture medium significantly enhanced cell growth, yielding a 9.44 ± 0.78-fold increase in total cell number by day 10 compared to day 1 (mean ± SEM) (Fig. 3d, e). In parallel, ALB secretion was markedly higher in Matrigel-supplemented cultures compared to controls without Matrigel (Fig. 3f). Additionally, direct thawing of cryopreserved HBs into Matrigel led to robust cellular outgrowth (Fig. 3g and Supplementary Fig. 3d), and ALB secretion correlated positively with increasing cell number (Fig. 3h). We further compared ALB secretion across different expansion conditions using ELISA (Supplementary Fig. 3e). By day 10 of culture, HBs maintained in SCF19 exhibited significantly higher ALB secretion compared to those cultured in alternative expansion media C and D, with the latter showing approximately 50% lower levels (Fig. 3i). These findings that demonstrate superior hepatic function and stability under SCF19 conditions were reproducible across multiple hPSC lines, as shown in Supplementary Figures 3f, 3g, and 3h.

In summary, HBs expanded in SCF19 not only retained hepatic progenitor characteristics but also demonstrated robust proliferative capacity following cryopreservation and single-cell seeding, supporting the suitability of this protocol for scalable cell manufacturing applications.

### Thyroid hormone and Wnt signaling promote hepatic maturation and zonation of hPSC-derived hepatocytes

In our previous study, we demonstrated that cyclic AMP (cAMP) treatment promoted hepatic maturation in 3D aggregates, as evidenced by the upregulation of multiple hepatic genes^33^.

However, even under these conditions, approximately 30% of the cell population retained expression of AFP-a marker of HBs that is not expressed in adult hepatocytes (Fig. 4a, b). During mammalian development, thyroid hormone has been implicated as a key regulator of organ maturation^34–38^. To determine whether thyroid hormone facilitates the maturation of hepatoblasts, we examined its effect on the differentiation of ALB⁺/AFP⁺ cells. To this end, monolayer cultures were maintained until day 33 without passaging, after which cells were reaggregated into 3D spheroids and cultured in DEX/OSM-containing medium for five days. On day 38, aggregates were treated with either triiodothyronine (T3) or cAMP for 18 days, followed by flow cytometric analysis of ALB and AFP expression. As shown in Figure 4a, treatment with T3 resulted in a significant reduction in AFP expression in 3D aggreagetes (Fig. 4a, b). RT-qPCR analysis confirmed these findings, revealing a complete reduction of AFP expression in the T3-treated population (Fig. 4c). Notably, AFP expression was lower in the cAMP- and T3-treated groups when compared to the untreated group (Fig. 4c).

**Figure 4.**
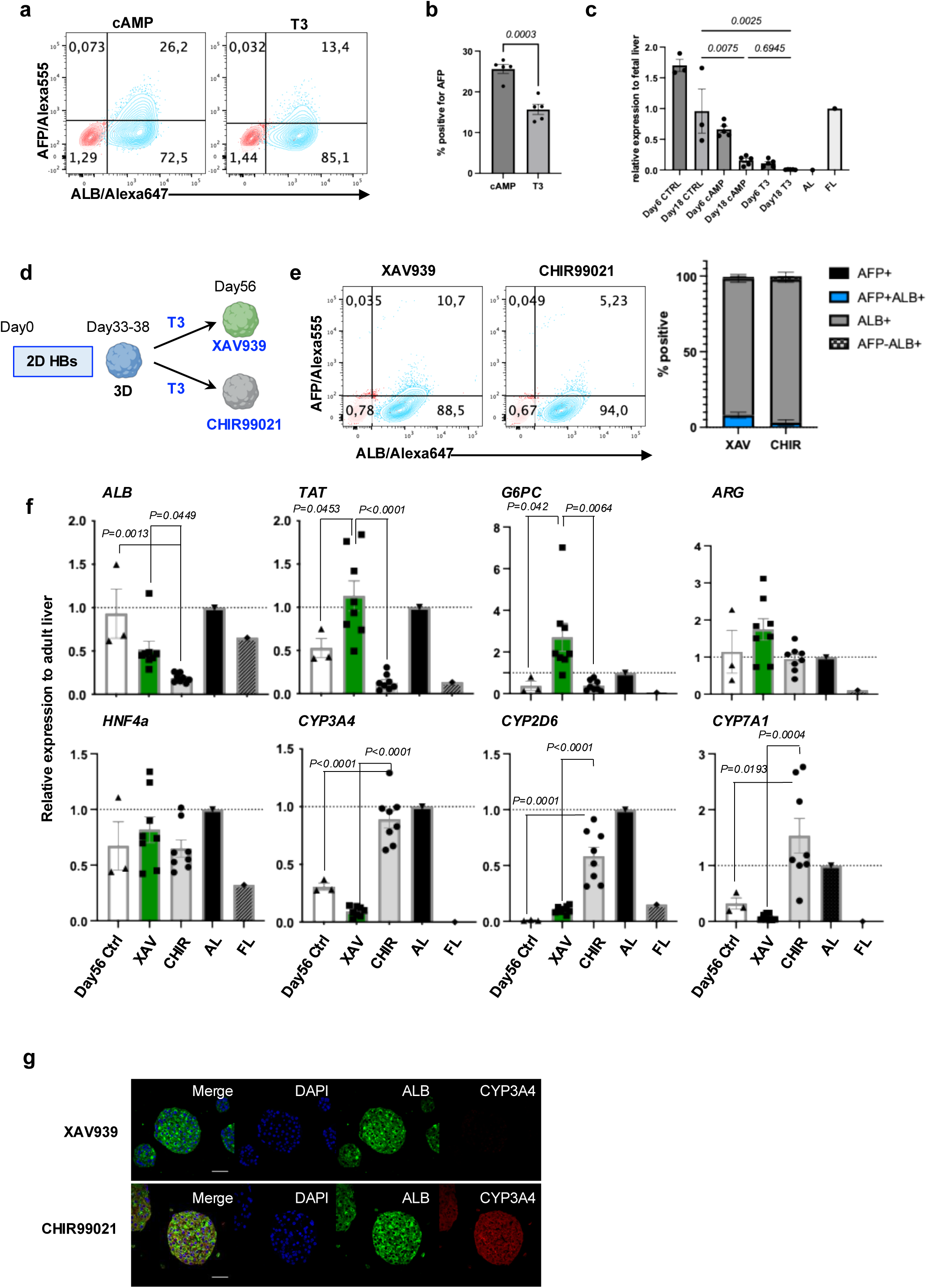
Thyroid hormone and Wnt signaling promote hepatic maturation and zonation of hPSC-derived hepatocytes. a) Flow cytometry analysis showing the population of ALB+ and AFP+ cells in cAMP or T3-treated hepatic aggregates. b) Quantification of the cell populations indicated in Fig. 4a. Two-tailed Student’s test (*n* = 5). c) RT-qPCR analysis of AFP expression in each population at day 6 and day 18 of H9-derived hepatic aggregates. AL adult liver, FL fetal liver. One-way ANOVA. DATA are represented as mean ± SEM (*n* = 3-5). d) Schematic differentiation of 3D hepatic aggregates. e) Flow cytometry analysis showing the population of ALB+ and AFP+ cells in XAV939 or CHIR99021-treated hepatic aggregates (left). Quantification of positive cells in each quadrant, indicated by colours (right) (*n* = 5). f) RT-qPCR analysis of the expression of the indicated genes in each population at day 56 of H9-derived hepatic aggregates under different culture conditions. AL adult liver, FL fetal liver. One-way ANOVA. DATA are represented as mean ± SEM (*n* = 3-8). g) Immunostaining analysis showing the proportion of ALB and CYP3A4 in H9-derived hepatic aggregates at day 56 treated with XAV939 or CHIR99021. Scale bar: 50µm.

Wnt/β-catenin signaling is a key regulatory pathway that governs hepatic zonation, a phenomenon in which hepatocytes exhibit distinct metabolic profiles based on their spatial location along the porto-central axis^39–45^. The liver comprises heterogeneous hepatocyte subpopulations, each with specialized functions demarcated by their proximity to the portal vein (zone 1), midlobular region (zone 2), or central vein (zone 3)^46,47^. Single-cell RNA sequencing analyses of murine livers have revealed that nearly 50% of hepatocyte-expressed genes display zonation-specific patterns, underpinning the metabolic and functional heterogeneity observed across lobular zones^48,49^. Pericentral hepatocytes (zone 3), positioned in a region of high Wnt/β-catenin signaling, are enriched for genes involved in xenobiotic metabolism, including the cytochrome P450 enzymes. In contrast, periportal hepatocytes (zone 1), which reside in a low Wnt signaling environment, preferentially express genes associated with gluconeogenesis, urea cycle metabolism, and mitochondrial β-oxidation^50,51^.

To recapitulate hepatic zonation *in vitro*, we developed a staged differentiation strategy using Wnt modulators in combination with thyroid hormone. 3D aggregates were subsequently treated for 18 days with T3 along with either the Wnt agonist CHIR99021 or the Wnt inhibitor XAV939 (Fig. 4d). Flow cytometric analysis revealed that nearly 90% of hepatocytes generated under both conditions expressed ALB without AFP, demonstrating efficient hepatic maturation (Figure 4e). RT-qPCR analysis revealed that the treatment with Wnt antagonists significantly upregulated periportal-associated genes such as *TAT* and *G6PC*, while Wnt activation promoted expression of pericentral markers including *CYP3A4*, *CYP2D6*, and *CYP7A1* (Fig. 4f). Immunostaining further confirmed a greater abundance of ALB^+^/ CYP3A4^+^ cells, indicative of zone 3-like hepatocytes, in CHIR-treated aggregates (Fig. 4g).

To validate the mechanistic involvement of thyroid hormone signaling, we examined the expression of thyroid hormone receptor (THR) isoforms. THRα and THRβ were expressed throughout the differentiation process, with THRβ—the liver-predominant isoform—being significantly upregulated upon aggregation compared to monolayer culture, suggesting increased sensitivity to T3 in 3D configurations (Supplementary Fig. 4a). Additionally, we tested the THRβ-selective agonist GC-1, which has been shown to stimulate hepatic and pancreatic cell proliferation in rodent models^52^. Treatment with GC-1 in 3D aggregates, in combination with Wnt modulators, resulted in marked downregulation of AFP, as confirmed by flow cytometric analysis (Supplementary Fig. 4b).

Together, these results demonstrate that T3 facilitates hepatic maturation more effectively than cAMP and that thyroid hormone signaling through T3 or the THRβ-specific agonist GC-1 can be leveraged to promote the maturation of hPSC-derived hepatocytes. Furthermore, these results suggest that spatially distinct hepatic functions can be established *in vitro* through the fine-tuned modulation of Wnt signaling.

### Single-cell RNA analyses of Wnt-zonated hPSC-derived hepatocytes

To elucidate the transcriptional heterogeneity and functional chracteristics of hPSC-derived zonated hepatocytes, we performed single-cell RNA sequencing (scRNA-seq) analysis. In the Wnt inhibitor (XAV939)-treated group, 324 individual cells were profiled, yielding transcriptomic data for 15,070 genes. In parallel, 424 cells were analyzed from the Wnt agonist (CHIR99021)-treated group. Clustering analysis identified transcriptionally distinct populations within each treatment group, designated as XAV-0 (Supplementary Data 1), XAV-1 (Supplementary Data 2), and XAV-2 (Supplementary Data 3) in XAV-reated geoup, and CHIR-0 (Supplementary Data 4), and CHIR-1 (Supplementary Data 5) in the CHIR-treated group, respectively (Fig. 5a).

**Figure 5.**
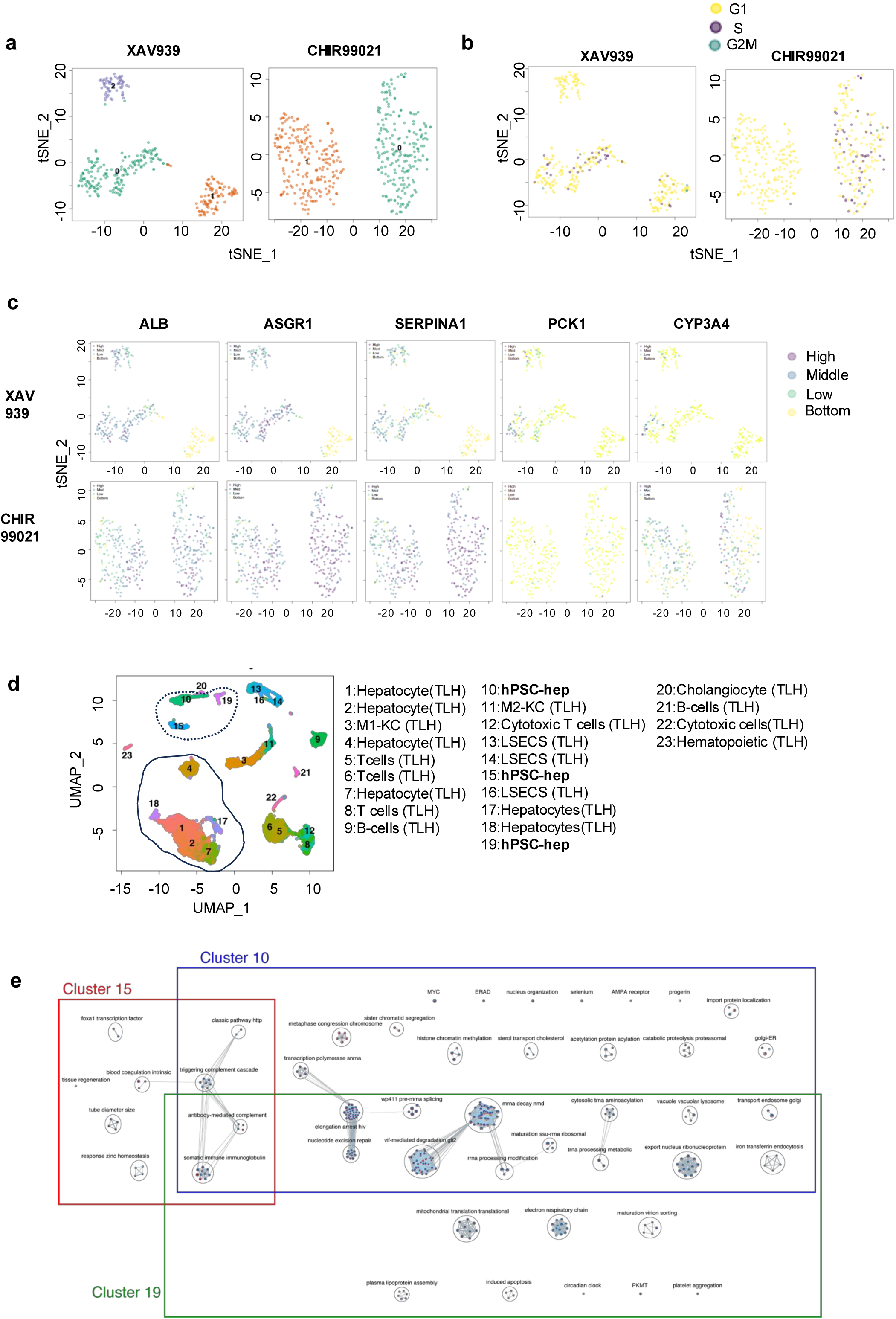
Global gene expression analysis reveals zonation features in hPSC-derived hepatocytes. a) tSNE plot of 3D hepatic aggregates treated with XAV939 or CHIR99021. b) tSNE plots showing cells in various cell cycles. c) tSNE plots displaying the expressions of selected hepatic markers. d) UMAP projection of integrated single-cell transcriptomic profiles from human adult liver cells and 3D hepatic aggregates derived from H9 hPSCs. Clusters corresponding to hPSC-derived hepatocytes are demarcated by dotted circles (Clusters 10, 15, and 19), whereas clusters representing primary human hepatocytes from adult liver tissue are enclosed with solid circles (Clusters 1, 2, 4, 7, 17, and 18). e) Pathway enrichment analysis comparing hPSC-derived hepatocyte clusters (Clusters 10, 15, and 19) to adult primary hepatocyte clusters (Clusters 1, 2, 4, 7, 17, and 18), highlighting signaling pathways significantly downregulated in hPSC-derived populations.

As shown in Figure 5b, cell cycle analysis revealed signatures of active proliferation across several clusters, characterized by distinct expression profiles associated with the G1, S, and G2/M phases. Notably, subpopulations within the XAV-2 (WNT-inhibited) and CHIR-0 (WNT-activated) clusters exhibited minimal expression of cell cycle–associated genes, indicating a predominantly quiescent or terminally differentiated cellular state. Expression of canonical hepatocyte marker genes, including *ALB*, *ASGR1*, and *SERPINA1* (*AAT*), was broadly detected across clusters, except in XAV-1, confirming successful hepatic lineage specification in most populations. Notably, *PCK1*, a hallmark gluconeogenic gene associated with periportal (zone 1) hepatocytes, was enriched in XAV-treated cells, whereas *CYP3A4*, a prototypical marker of pericentral (zone 3) hepatocytes, was preferentially expressed in CHIR-treated aggregates (Fig. 5c).

To further contextualize these *in vitro* phenotypes, we integrated our scRNA-seq data with a previously established human liver single-cell atlas^50^. Uniform Manifold Approximation and Projection (UMAP) visualization of integrated scRNA-seq data revealed transcriptomic similarities between hPSC-derived hepatocyte clusters and primary human hepatocytes (Fig. 5d). Clusters XAV-0 (Wnt-inhibited) and CHIR-0 (Wnt-activated) co-localized within a composite Cluster 10, suggesting a transcriptional identity resembling periportal and midzonal hepatocytes. In contrast, Cluster 15, comprising cells from both XAV-2 and CHIR-1, was predominantly populated by CHIR-1-treated cells (78.5%; 212/270) (Supplementary Fig. 5a, b). Differential gene expression analysis between Clusters 10 and 15 demonstrated distinct zonation-like transcriptomic signatures. Pathway enrichment analysis showed that Cluster 15 was significantly enriched for genes involved in bile acid biosynthesis and xenobiotic metabolism, consistent with a pericentral (zone 3) identity and reflecting the impact of Wnt activation on hepatic zonation (Supplementary Fig. 6).

Intriguingly, XAV-1 formed a transcriptionally unique cluster (Cluster 19) with limited overlap with canonical hepatocyte zones. Instead, Cluster 19 appeared to represent an intermediate or divergent cell state, possibly of progenitor-like or non-parenchymal origin. Correlation analysis of top marker genes in each cluster against zonated regions in the Human Liver Atlas indicated that Cluster 15 closely resembled midzonal to zone 3 hepatocytes, but did not correlate with layer 1 cells immediately adjacent to the central vein (Supplementary Fig. 7, Supplementary Data 6). Cluster 10 showed strong similarity to periportal and midzonal hepatocytes, supporting its classification as a zone 1-like population. Notably, Cluster 19, entirely derived from the XAV-treated group, was enriched for transcripts associated with cholangiocytes and M2 macrophages, and exhibited partial transcriptional resemblance to hepatocytes located near both portal and central veins. These findings suggest that Wnt inhibition can induce spatial and lineage heterogeneity in 3D hepatic cultures, potentially giving rise to subpopulations with non-parenchymal or hybrid identities due to residual endogenous Wnt signaling.

To further evaluate the degree of transcriptomic convergence with mature human hepatocytes, comparative pathway analysis was conducted, revealing several signaling cascades that were underrepresented in hPSC-derived hepatocytes (hPSC-HEPs), particularly in Cluster 15, which otherwise displayed the highest similarity to primary adult hepatocytes. Despite this resemblance, Cluster 15 lacked expression of several key hepatic programs, including pathways regulated by the transcription factor *FOXA1*, as well as genes involved in coagulation, tissue regeneration, vascular development, and zinc homeostasis (Fig. 5e)^53^. A more detailed comparison among Clusters 10, 15, and 19 revealed a consistent absence of numerous liver-enriched genes in hPSC-derived cells compared to their *in vivo* counterparts (Supplementary Data 7).

Importantly, none of the hPSC-derived clusters expressed acute-phase proteins such as *HP* (haptoglobin) or *FGA/B* (fibrinogen alpha/beta chains), nor genes related to metal ion metabolism, including *MT1X*, *MT1G*, and *MT2A* (Supplementary Data 7). In contrast, genes enriched in hPSC-derived hepatocytes but absent in mature liver tissue included bile duct-associated transcription factors such as *SOX4* and *LGALS*. Notably, hPSC-hepatocyte cluster 10 and 19 populations showed significantly elevated expression of genes within several signaling patways (Supplemetary Fig. 5c), relative to primary adult hepatocytes. While these transcripts were not detected in Cluster 15, they were highly expressed in Clusters 10 and 19, along with *SOX9* and *GPC3*, the latter being a known marker of proliferative or immature hepatocytes (Supplementary Data 7)^54–56^. These expression patterns indicate that Clusters 10 and 19 retain a residual progenitor-like or biliary lineage signature, reflecting incomplete differentiation or cellular heterogeneity within the hPSC-derived populations.

Taken together, our data suggest that while specific subpopulations of hPSC-derived hepatocytes, particularly those in Cluster 15, can approximate the transcriptional identity of mature hepatocytes, other populations retain progenitor-like or non-parenchymal traits. This highlights the need for further optimization of differentiation protocols to fully recapitulate the transcriptional complexity and functional repertoire of primary human hepatocytes.

### Generation of mature hepatocytes and cliated cholangiocyte from cryopreserved hepatoblasts

Regenerative medicine-based therapies depend on the ability to efficiently and reproducibly generate sufficient numbers of the target cell of interest. While the advances described in this study establish methods to generate relatively mature hPSC-derived hepatocytes in 3D aggregates, the total cell yield remains suboptimal. To address this limitation, we set out to design a protocol that would promote a more efficient generation of mature hepatic cells from cryopreserved HBs. For this approach, we tested the effect of different signaling pathway agonists and antagonists in a 2D culture format. From these analyses, we found that modulation of the Wnt pathway by treatment with either CHIR99021 or XAV939 together with inhibition of both TGF-β (SB431542) and Notch signaling suppressed cholangiocytic lineage commitment and promoted hepatocytic specification. (Fig. 6a). Flow cytometry and immunohistochemistry revealed a gradual reduction in AFP expression, indicating progressive maturation over a 24-day period (Fig. 6b, c). Notably, supplementation with low concentrations of FGF19 led to a 30–50% increase in the yield of mature ALB^+^/AFP^-^ hepatocytes by day 24, suggesting enhanced expansion potential during terminal differentiation (Fig. 6d). At day 12 of maturation, the cells were aggregated into 3D spheroids and further matured for an additional 12 days in floating culture, in the presence of T3 and either a Wnt activator or inhibitor (Fig. 6e and Supplementary Fig. 8a). qPCR analysis revealed significantly higher expression of the zone 3 marker *CYP3A4* in CHIR-treated cells compared to XAV-treated cells, even following the combined 2D and 3D maturation process (Supplementary Fig. 8b).

**Figure 6.**
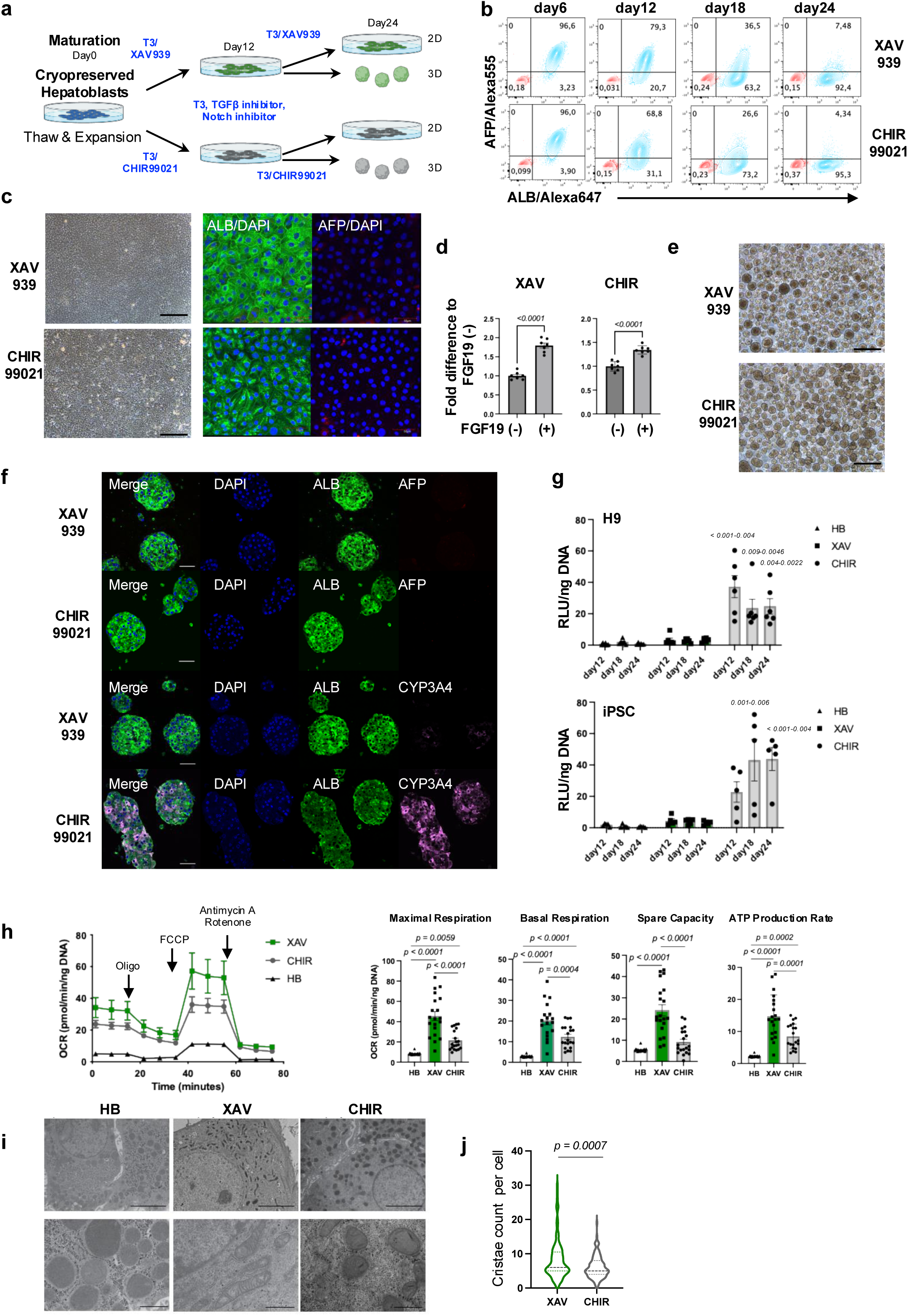
Generation and functional characterization of mature hepatocytes from cryopreserved HBs a) Schematic differentiation protocol of hPSC-derived hepatocytes from cryopreserved hepatoblasts. b) Flow cytometry analysis showing the population of ALB+ and AFP+ cells with XAV939 or CHIR99021-treated hepatic cells over day-24 period. c) Microscopic (left) and Immunostaining (right) analysis, showing the proportion of ALB (green), and AFP (red) with XAV939 or CHIR99021-treated hepatic cells. Scale bar: 200µm (left), 50µm (right). d) Quantification of fold change in cell numbers with XAV939 or CHIR99021-treated hepatic cells treated in the presence or absence of FGF19. Student’s t-test. DATA are represented as mean ± SEM (*n* = 7). e) Microscopic analysis showing hepatic aggregates treated with XAV939 or CHIR99021. Scale bar: 200µm. f) Immunostaining analysis showing the proportion of ALB/AFP (top) and ALB/CYP3A4 (bottom) in H9-derived hepatic aggregates treated with XAV939 or CHIR99021. Scale bar: 50µm. g) CYP3A4 activity in H9- and iPSC-derived hepatic aggregates at day12, 18 and 24 of maturation treated with XAV939 or CHIR99021. CHIR-treated groups exhibited higher CYP3A4 compared to both HB and XAV-treated groups. DATA are represented as mean ± SEM. One-way ANOVA. Statistical significance compared to HB and XAV groups is indicated in Figure (H9: *n* = 6, iPS *n* = 5). h) Representative kinetics of the oxygen consumption rate (OCR) in H9-derived day 24 mature hepatic aggregates cultured in indicated conditions (left). Comparison of each parameter in the day 24 mature hepatic aggregates (right). One-way ANOVA. DATA are represented as mean ± SEM from 4 technical replicates of 5 biologically independent experiments. i) Representative transmission electron microscope images of mitochondria in different treated hepatic aggregates under the indicated conditions. Scale bar: 5µm (top), 1µm (bottom). j) Quantification of cristae found in mitochondria under the indicated conditions. Data are from 3 biologically independent experiments, Student’s t-test. DATA are represented as mean ± SEM.

Consistently, immunofluorescence analysis demonstrated robust CYP3A4 protein expression, particularly within organoids treated with the Wnt activator CHIR99021 (Fig. 6f). In addition, immunohistochemical staining confirmed the zonal distribution of other key hepatocyte functional markers, further supporting the successful induction of spatially distinct hepatic phenotypes through Wnt signaling modulation (Supplementary Fig. 8c). Functional assays further confirmed that CYP3A4 metabolic activity was differentially modulated by Wnt signaling, with Wnt activation significantly enhancing drug metabolism capacity compared to Wnt inhibition (Fig. 6g). Notably, CYP3A4 activity in CHIR-treated organoids was sustained for over 12 days, suggesting a more stable induction of zone 3-like hepatic function under Wnt-activated conditions. Similarly, during the maturation of cryopreserved iPSC-derived hepatoblasts, higher CYP3A4 metabolite activity was consistently observed under Wnt-active conditions relative to Wnt-inhibited conditions, further supporting the role of Wnt signaling in promoting and maintaining pericentral hepatocyte functionality. This indicates that our maturation condition enables the robust and reproducible generation of zone 3-like hepatocytes from expandable progenitor cell banks. When CYP3A4 activity was assessed in hepatocytes isolated from humanized liver mice^57,58^ and cultured under T3-XAV or T3-CHIR, enzymatic activity peaked at day 6 in the CHIR condition and declined thereafter (Supplementary Fig. 9a). This transient activity likely reflects the inherent instability and fragility of mature human hepatocytes under *in vitro* conditions.

Moreover, we evaluated the potential of cryopreserved HBs expanded under SCF19 conditions to differentiate into cholangiocytes. Using our previously published protocol for cholangiocyte induction, frozen-thawed HBs were efficiently directed to differentiate into primary cilia-positive cholangiocytes in monolayer culture^27^. These cells expressed key bile duct-specific markers, including *CFTR,* and demonstrated functional capacity by forming 3D bile duct-like cystic organoids in suspension culture, confirming their biliary lineage commitment and morphogenetic potential (Supplementary Fig. 10a-d).

### Differential mitochondrial function in hPSC hepatocytes following Wnt modification

Mitochondrial function is a critical aspect of hepatic metabolism, and zone-specific metabolic differences have been well documented within the liver lobule^59^. To assess mitochondrial activity in hPSC-derived hepatocytes exhibiting zonal phenotypes, we employed the Seahorse XF Analyzer to measure oxygen consumption rate (OCR) and ATP production rate during OCR in 3D aggregates generated under Wnt-inhibited and Wnt-activated conditions. Hepatocytes treated with the Wnt inhibitor XAV939 exhibited significantly higher basal respiration, maximal respiration, spare capacity, and ATP production rate compared to those treated with the Wnt activator CHIR99021 (Fig. 6h). These results indicate a metabolic phenotype resembling that of zone 1 hepatocytes, which are characterized by elevated oxidative phosphorylation due to their proximity to the portal vein. Supporting these observations, qPCR analysis revealed differential expression of key mitochondrial and metabolic genes in day 24 mature 3D aggregates, as well as in mature single cells embedded in collagen gel following day 12 monolayer-based maturation, in accordance with our established protocol (Supplementary Fig. 11a, b). Both 3D aggregates and single cells embedded in collagen gels exhibited maturation property, as evidenced by reduced AFP expression detected by flow cytometry analysis (Supplementary Fig. 11c). *CPT1A*, a rate-limiting enzyme in fatty acid β-oxidation, along with *PGC1a,* was significantly upregulated under Wnt-inhibited conditions. Similarly, *MFN2* and *OPA1*, genes involved in mitochondrial fusion and turnover, exhibited elevated expression under Wnt inhibition conditions, suggesting enhanced mitochondrial dynamics. In contrast, the glycolysis associated gene *Glut2* was highly expressed in Wnt-activated cells. Although zone 3 hepatocytes are predominantly associated with glycolytic activity, it is notable that *PFKL*— another key regulatory enzyme in glycolysis—was also upregulated under Wnt-inhibited conditions (Supplementary Fig. 11d). This suggests that, despite enhanced oxidative metabolism in XAV-treated cells, a degree of glycolytic activity is retained, indicative of a mixed metabolic phenotype.

Furthermore, comparative analysis with transcriptomic data from primary human hepatocytes corroborated these findings by revealing enrichment of oxidative metabolic pathways under Wnt-inhibited conditions (Supplementary Fig. 9b, c), consistent with elevated fatty acid oxidation and mitochondrial energy output. Collectively, these results indicate that Wnt signaling not only governs zonal identity in hPSC-derived hepatocytes but also drives zone-specific metabolic reprogramming reflective of physiological hepatic function.

Transmission electron microscopy (TEM) provided ultrastructural validation, revealing that hepatic progenitor cells possessed mitochondria with sparse or absent cristae, whereas differentiated hepatocytes, particularly under XAV treatment, displayed dense cristae networks and increased inner membrane surface area (Fig. 6i, j). These morphological changes, aligned with Seahorse functional data, support the conclusion that Wnt-inhibited conditions promote mitochondrial maturation and respiratory competence, contributing to a more physiologically relevant zone 1-like hepatic phenotype.

### *In vivo* potential of hPSC-derived hepatocytes

Finally, to assess the functional capacity of hPSC-derived hepatocytes *in vivo*, we transplanted the cells into immunocompromised mice and evaluated graft viability and hepatic function. 3D hepatic aggregates generated from HBs under different maturation conditions— XAV939 (Wnt inhibitor), CHIR99021 (Wnt activator), and a 1:1 combination of XAV and CHIR-treated cells—were transplanted beneath the kidney capsule of immunodeficient mice. The HB-transplanted group exhibited negligible human albumin secretion at 4 weeks post-transplantation, and histological analysis revealed a predominant conversion of cells into ALB^-^, CK7 ^+^ cholangiocyte-like cells, indicating lineage diversion. In contrast, the XAV-treated and CHIR-treated groups demonstrated measurable levels of human albumin secretion (298.4 ± 84.6 ng/mL and 250. 4 ± 110.568ng/mL, respectively) at 4 weeks, although albumin production declined progressively over time. Notably, the group receiving a mixture of XAV- and CHIR-treated cells showed the highest levels of albumin secretion (499.8 ± 65.8ng/mL) than mice transplanted with either population alone (Supplementary Fig. 12a). Immunostaining of the grafts revealed a greater abundance of ALB^+^ hepatocyte-like cells in the population generated from the mixed zonated cells (Supplementary Fig. 12b). However, CYP3A4 expression, a marker of zone 3 hepatocytes, was not detected in any of the transplanted groups by immunohistochemistry at this site.

To further assess hepatic engraftment and functional integration within a physiologically relevant hepatic niche, we transplanted 1 million cells from each group (HB, XAV-treated, and CHIR-treated) into the spleen of TK-NOG mice^57,58^. These mice carry a genetically engineered HSV-TK transgene driven by the mouse albumin promoter, enabling selective ablation of host hepatocytes upon administration of ganciclovir (GCV), which was administered prior to cell transplantation. At 12 weeks post-transplantation, human albumin was undetectable in the HB group, while the XAV-treated group showed measurable levels of human albumin (1048.267 ± 521.090 ng/mL; *n=*15) (Supplemtary Fig. 12c). Immunohistochemical analysis at 16 weeks confirmed the presence of small clusters of ALB^+^, human KU80^+^ cells within the liver parenchyma of recipient mice. In contrast, the CHIR-treated group exhibited markedly lower albumin production, indicating a limited contribution to intrahepatic repopulation. Compared to the control group transplanted with primary human hepatocytes (PH), the engraftment efficiency of hPSC-derived hepatocytes remained low (<5%), however, immunostaining revealed the presence of CYP3A4^+^ cells among the engrafted population (Supplementary Fig. 12d).

### 3D bioprinted liver tissue generation using scalable 3D-hep platform

To further leverage our scalable culture platform, we explored the fabrication of structured liver tissue constructs using an extrusion-based 3D bioprinter (Fig. 7a)^60–63^. HBs, preserved via our cryopresercation protocol, were thawed and directly employed to generate HB-, XAV-treated, and CHIR-treated organoids. Zone specific maturation was first induced under monolayer conditions until day 12, after which cells were reaggregated and maintained as 3D organoids in suspension culture for an additional four days prior to biopringing. A total of 1 mL of bioink, composed of laminin and alginate, was combined with 20 million cells from each group (HBs, XAV-treated, and CHIR-treated organoids). Printing parameters, including extrusion speed and pressure, were optimized to maintain structural fidelity and prevent deformation. Under optimal conditions (extrusion speed: 3 mm/sec; pressure: 3–5 psi), we successfully fabricated three-layered hepatic tissue constructs measuring 10 mm×10 mm×1 mm within a single well of a standard 6-well plate (Fig. 7b). Following bioprinting, the constructs were cultured under lineage-specific conditions designed to preserve the zonal identity of the organoids. Cell viability was assessed using Live/Dead staining, which confirmed the survival of embedded organoids within the bioink matrix (Supplementary Fig. 13). Albumin secretion from the grid tissue was measured, and even at 8 days post-print, measurable albumin secretion was observed, indicating preserved hepatic function (Figure 7c). At 8 days post-printing, total RNA was extracted from the grid-like constructs and analyzed by qPCR. Expression of *ALB*, *AFP*, and *CYP3A4* was maintained, reflecting preservation of hepatic lineage commitment and functional identity (Fig. 7d).

**Figure 7.**
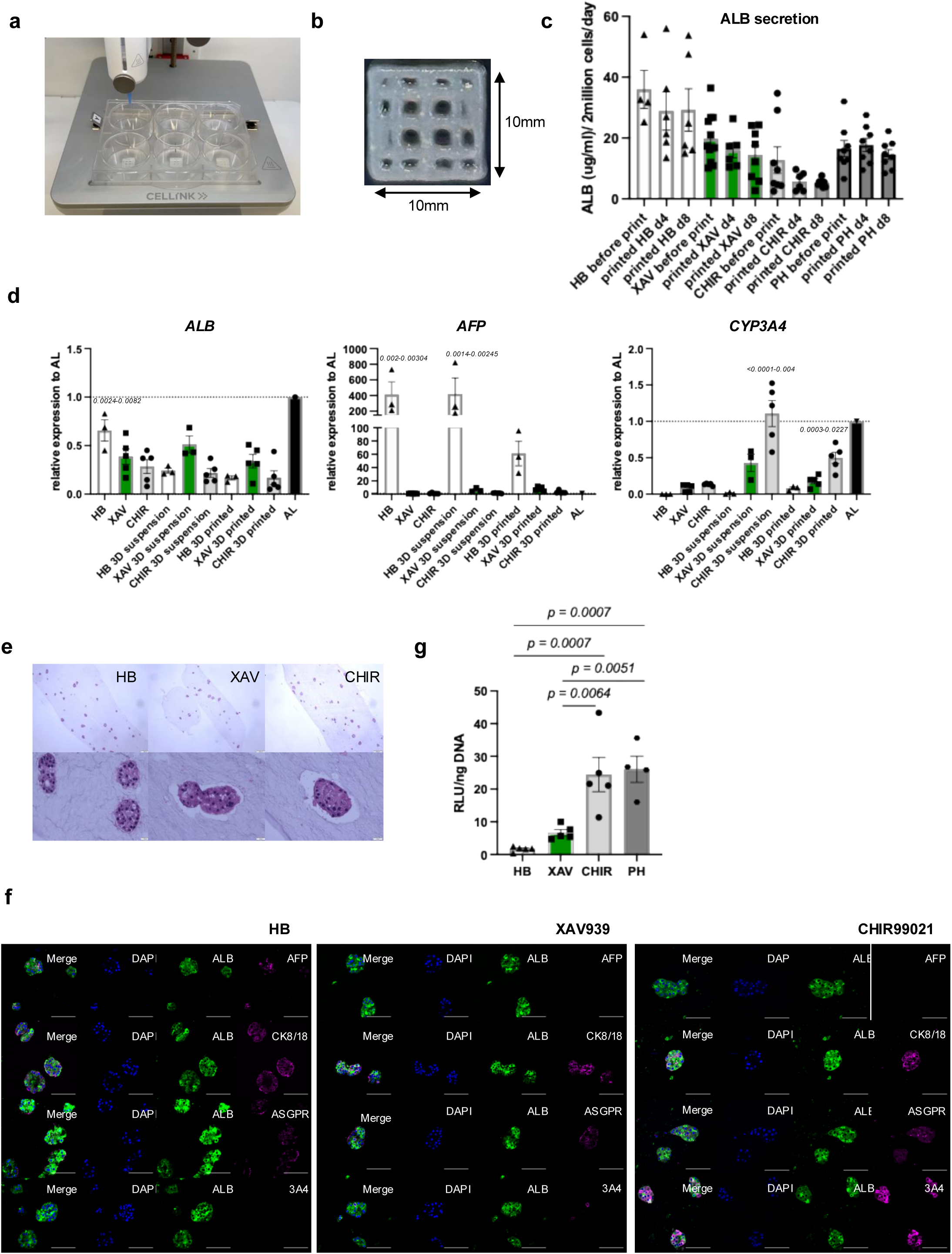
3D bioprinted hPSC-derived hepatic aggregates exhibited functional properties. a) 3D hepatic aggregates were printed by CELLINK BIOX 3D Bioprinter. b) Image showing a grid structure in which cells were bioprinted. c) Quantification of ALB secretion before bioprinting (3D) and after bioprinting at day4 and day8 under different conditions (*n* = 4-12). d) RT-qPCR analysis of indicated gene expression in monolayer, 3D suspension, and bioprinted cultures under different conditions. AL adult liver, One-way ANOVA. DATA are represented as mean ± SEM. Statistical significance compared to other groups is indicated in the Figure (*n* = 3-5). e) Representative microscopic image showing H and E staining of 3D printed hepatic aggregates Scale bar: 200 µm (top), 20µm (bottom). f) Representative confocal image showing the proportion of co-staining with ALB, and AFP (top), CK8/18 (second row from the top), ASGPR (third from the top), CYP3A4 (bottom) in HB, XAV989 or CHIR98014-treated hepatic cells. Scale bars: 100µm. g) CYP3A4 activity in bioprinted hepatic aggregates at day8 post-printing.

Notably, *CYP3A4* expression remained elevated in CHIR-treated (Zone 3–like) constructs, consistent with the original 3D organoid phenotype prior to printing. Histological analysis revealed well-preserved organoid morphology within the bioink matrix (Fig. 7e). Immunostaining showed that HB-derived constructs exhibited ALB/AFP double-positive cells, while XAV-treated tissues contained predominantly AFP^-^, ALB^+^, and ASGR1^+^ hepatocytes, yet lacked detectable CYP3A4 expression. In contrast, CHIR-treated constructs displayed numerous ALB^+^/CYP3A4^+^ organoids, indicative of a Zone 3–like phenotype (Fig. 7f). These results were corroborated by functional assays, wherein CYP3A4 metabolic activity in CHIR-treated bioprinted tissues was significantly higher than in other groups and comparable to printed constructs prepared using primary human hepatocytes (Fig. 7g).

Collectively, these findings demonstrate that hepatic organoids generated from hPSC-derived expandable hepatic progenitors can be successfully embedded and maintained within 3D bioprinted constructs while preserving zonal metabolic characteristics. The ability to modulate Wnt signaling enables the generation of distinct zone-specific hepatocyte subtypes within a structurally stable and scalable engineered platform. This system offers a promising foundation for the development of functional, transplantable liver tissue as well as physiologically relevant models for drug screening and disease modeling.

## Discussion

Despite substantial progress in hPSC-based liver differentiation, the scalable generation of functionally mature and zonally organized hepatocytes remains a critical barrier to clinical translation. While bioreactor-based suspension systems have demonstrated success in scaling undifferentiated hPSCs, their advantage is largely restricted to the pluripotent stage^64–66^. Once cells undergo lineage commitment, particularly toward definitive endoderm or hepatic lineages, the expansion potential rapidly declines. These systems lack the environmental precision necessary for lineage-specific amplification or maturation, leading to heterogeneous and often functionally compromised cell populations.

In contrast, our study establishes a chemically defined and reproducible platform for the post-pluripotent, scalable expansion of hepatic progenitor cells (HPCs) derived from hPSCs, representing a key advancement over existing methods that are largely restricted to the undifferentiated stage. While our previously reported protocol enabled the induction of approximately 7–10 HPCs per hPSC^27^, the current study demonstrates that each HPC can undergo 6- to 10-fold expansion per passage. When combined with cryopreservation and serial re-expansion, this system theoretically enables the generation of nearly 100 HPCs per starting hPSC within a single passage, with continued 10-fold expansion achievable in subsequent cycles. Notably, this entire expansion process occurs in a monolayer culture format under serum-free, chemically defined conditions, eliminating the need for ECM-dependent cystic organoids, which often suffer from limited scalability, high cost, and batch variability. By circumventing these constraints, our platform provides a highly cost-effective and reproducible solution for generating lineage-committed hepatic cells at clinically relevant scales.

Unlike conventional protocols for hepatic organoid generation or hepatic progenitor expansion, which rely heavily on mitogens such as HGF and EGF or conditioned media to supply Wnt ligands, our approach sustains hepatic progenitor identity and bipotency under fully defined conditions. Critically, it minimizes off-target differentiation into CK19⁺ cholangiocyte-like cells that lack functional characteristics, while preserving the capacity to generate primary cilia-bearing, functionally competent cholangiocytes under directed differentiation. This versatility allows for the serial expansion, cryopreservation, and on-demand maturation of HPCs into either high-purity hepatocytes or cholangiocytes, offering a flexible and scalable platform compatible with both research applications and preclinical manufacturing pipelines.

Our findings further highlight the functional competence of hepatocytes generated from this platform. Consistent with a prior report indicating that thyroid hormone promotes hepatic maturation via activation of the CYP3A4 promoter thourgh T3 and thyroid hormone receptor analogues^20^, we achieved an efficient transition from AFP to ALB, a hallmark of hepatocyte maturation. Importantly, WNT pathway modulation enabled the induction of hepatic zonation *in vitro*, generating zone 3-like hepatocytes with CHIR99021 and zone 1-like profiles with XAV939. Single-cell transcriptomic analysis confirmed that these populations exhibit transcriptional signatures mirroring the metabolic gradients observed *in vivo* across the liver lobule. Interestingly, XAV-treated organoids exhibited a subpopulation resembling pericentral layer 1 cells despite Wnt inhibition, suggesting the persistence of endogenous WNT niches or spatial self-organization within 3D cultures.

While hPSC-derived hepatocytes recapitulated many features of mature liver tissue, transcriptomic comparison with primary adult hepatocytes revealed deficiencies in pathways linked to metal ion metabolism (e.g., zinc), acute-phase response (e.g., HP, FGA/B), and complement signaling. These deficits underscore the inherent limitations of monoculture systems, which lack the cellular cross-talk provided by hepatic stellate cells (HSCs), sinusoidal endothelial cells, and immune cell types. Incorporation of such cellular niches or engineering synthetic equivalents may further close this functional gap.

Importantly, transplantation studies validated the functional potential of our cells *in vivo*. Under the renal capsule, co-transplantation of CHIR- and XAV-treated hepatocytes enhanced albumin secretion, suggesting cooperative interaction between zone-specific subtypes, athough CYP3A4 was not expressed at this ectopic site. In contrast, intrahepatic transplantation into humanized TKNOG mice led to stable engraftment, sustained albumin secretion, and upregulation of CYP3A4 in XAV-treated cells grafts, confirming environmental reprogramming and phenotypic plasticity toward zone 3 function.

Finally, our system is compatible with tissue engineering platforms. Using extrusion-based 3D bioprinting, we fabricated structured hepatic tissues incorporating zonally specified organoids. CHIR-derived constructs maintained CYP3A4 expression and enzymatic activity equivalent to that of primary hepatocyte-printed tissue, demonstrating the potential of our platform for use in drug screening, disease modeling, and regenerative applications. Many 3D liver organoid models that rely on undefined ECMs such as Matrigel hinder scalability, batch consistency, and clinical translation. In contrast, our strategy enables expansion and maturation in scaffold-free conditions up to the progenitor stage, with flexibility in matrix selection for final tissue fabrication.

In summary, we present a robust, flexible, and scalable strategy for the generation, banking, and maturation of hPSC-derived hepatic progenitor cells into zonally patterned hepatocytes. By decoupling expansion from pluripotency and enabling downstream tissue engineering integration, this platform addresses long-standing limitations in hepatocyte manufacturing and offers a powerful tool for both clinical and industrial translation.

## Methods

### Differentiation of human ES and iPS cells into hepatoblasts in monolayer culture

Human embryonic stem cell (hESCs; H9/WA009 and GCaMP3) and induced pluripotent stem cells (iPSCs; WT01 and CF02a) were cultured on irradiated mouse embryonic fibroblasts in hESC medium. Differentiation of hPSCs into hepatoblasts was performed as previously described with minor modifications^27^. Briefly, prior to endoderm induction, hPSCs were seeded onto 2.5% Matrigel (Corning: 354230)-coated 12-well plates at a density of 100,000–150,000 cells per well. To initiate definitive endoderm differentiation, cells were cultured in RPMI 1640 medium (Thermo Fisher: 11875093) supplemented with glutamine (2 mM, Thermo Fisher: 25030081), 4.5X10^-4^M monothioglycerol (MTG; Sigma: M6145), activin A (100ng/ml, R&D: 388-AC/CF), and CHIR99021 (2µM, TOCRIS: 4423). On day 1, CHIR99021 was reduced to 1uM, and cells were maintained for two additional days in RPMI medium containing glutamine, ascorbic acid (50µg/ml, Sigma: A4544), monothioglycerol, basic fibroblast growth factor (bFGF, 5ng/ml, R&D: 233-FB), and activin A (100ng/ml) followed by four days in serum-free differentiation (SFD) medium with the same supplements. Media changes were performed every two days. On day 9, definitive endoderm identity was confirmed via flow cytometry for CXCR4 and cKIT expression. Cells were then directed toward hepatic specification using low-glucose DMEM (Thermo Fisher: 11885084) supplemented with bFGF (40ng/ml), Bone Morphogenic Protein 4 (BMP4, 50ng/ml, R&D: 314-BP), 1% B27 (v/v), (Thermo Fisher: 17504044), ascorbic acid, glutamine, and MTG. Media were changed every two days from days 9 to 15. For hepatoblast maturation, cells were cultured for eight days in a 3:1 mixture of low glucose DMEM (Thermo Fisher: 11885084) and Ham’s F12 (Corning: 10080CV), supplemented with 0.1% BSA, 1% B27 (v/v), ascorbic acid, glutamine, MTG, hepatocyte growth factor (HGF, 20ng/ml, R&D: 294-HGN), dexamethasone (Dex, 40ng/ml, BioShop: DEX002), and Oncostatin M (OSM; 20ng/ml, R&D: 295-OM/CF). From day 15 to day 23, 1 μM CHIR99021 was included for hPSC cell-derived hepatoblasts differentiation. All staged of differentiation-including endoderm induction, hepatic specification, and hepatoblasts maturation-were carried out under hypoxic conditions (5% CO2, 5% O2, 90% N2) through day 23. Cells were then transitioned to ambient oxygen conditions and maintained for an additional six days in a 3:1 mixture of high-glucose DMEM (Thermo Fisher: 11995065) and Ham’s F12, supplemented with 0.1% BSA, 1% B27 (v/v), ascorbic acid, glutamine, MTG, HGF (20ng/ml), Dex (40ng/ml), and OSM (20ng/ml).

### Expansion and cryopreservation of hPSC-derived hepatoblasts

After confirming cells were over 85% positive for ALB and AFP, with less than 1% of double-negative cells on day 29, cells were dissociated with TrypLE and re-seeded at a density of 100,000 cells per well onto pre-coated 2.5% Matrigel 12-well plates. The culture medium consisted of DMEM/F12 (Corning: 10092CV), supplemented with 0.1% BSA, 1% B27, 1% ITS (WISENT: 315082QL), 1% CDL (v/v), (Thermo Fisher: 11905031), ascorbic acid, glutamine, MTG, Dex (40ng/ml), 6uM SB (TOCIRS: 1614), 1uM CHIR99021 and FGF19 (50ng/ml, Prospec: CYT-700-3) (refered to **SCF medium**). ROCK inhibitor Y-27632 (RI) (10μM, TOCRIS: 1254) was added only on the first day. The medium was replaced every two days until the cells reached confluence. Once confluent, cells were passaged using the same protocol. In parallel, proliferated hepatoblasts were cryopreserved using a cryopreservation medium containing dimethyl sulfoxide (Sigma DMSO).

### Generation of 3D hepatic progenitor organoid in suspension culture and Matrigel

Cryopreserved hPSC-derived hepatoblasts (HBs) were seeded directly into Elplasia 96-well plates (Corning 4442), 6-cm dishes, or 10-cm dishes. Various seeding densities were evaluated, as described in the Results section. Cells were cultured in SCF19 medium, as previously described. When used, Matrigel was added at a final concentration of 0.75% (v/v). ROCK inhibitor (RI) and DNase were added on the first day of culture. In 96-well microcavity plates, the formation of cell aggregates was monitored using the Cell3iMager system (SCREEN Holdings). For cultures in 6-cm and 10-cm dishes, 3D aggregates were maintained on an orbital shaker within an ambient CO₂ incubator throughout the culture period.

### Maturation of 3D hepatic organoids and zonal specification from hPSC-derived hepatic progenitor cells

Human pluripotent stem cells (hPSCs: H9) were differentiated into hepatic progenitor cells over a 29-day period as previously described. For the initial maturation phase, day 29 hepatic progenitors were cultured for an additional 4 days in DMEM/F12 medium (Corning, 10092CV) supplemented with 0.1% bovine serum albumin (BSA), 1% insulin-transferrin-selenium (ITS; WISENT, 315082QL), ascorbic acid, L-glutamine, monothioglycerol (MTG), 40 ng/mL dexamethasone (Dex), and 2 ng/mL Oncostatin M (OSM; Thermo Fisher, 11905031).

On day 33, cells were enzymatically dissociated and transferred to ultra-low attachment 10 cm culture dishes at a seeding density of 5–15 × 10⁶ cells per dish to promote three-dimensional (3D) aggregation. Aggregates were maintained in DMEM/F12 supplemented with 0.1% BSA, 1% B27, 1% ITS, 1% chemically defined lipid concentrate (CDL; Thermo Fisher, 11905031), ascorbic acid, L-glutamine, MTG, and 40 ng/mL Dex. The ROCK inhibitor Y-27632 (10 μM; TOCRIS, 1254) was included only on day 33 to support aggregate formation. Fresh medium was replaced on days 34 and 36 to facilitate continued maturation. Beginning on day 38, aggregates were cultured for an additional 18 days in the same basal medium supplemented with either 1 mM 8-bromoadenosine 3′,5′-cyclic monophosphate (8-Br-cAMP) or 40 nM triiodothyronine (T3; Sigma-Aldrich) or 50nM GC 1 (Tocris, 4554/10) to enhance hepatic functional maturation.

For metabolic zonation, 2 μM XAV939 (TOCRIS, 3748) was added to induce Zone 1 (Z1)-like identity, whereas 1 μM CHIR99021 was added to promote Zone 3 (Z3) specification. All procedures were performed under ambient atmospheric conditions. 3D aggregates were maintained on the orbital shaker at 60 rpm speed. Control cells were treated with 0.33ul/ml of DMSO. Culture medium was refreshed every two days to ensure optimal nutrient supply and waste removal.

### Cholangiocyte differentiation from hPSC-derived cryopreserved hepatoblasts

Cholangiocyte differentiation was perfomed as previously described^27^. Cryopreserved hPSC (H9)-derived hepatoblasts were thawed, dissociated using TrypLE, and plated onto the irradiated OP9j cells. The cells were cultured in DMEM/Ham’s F12 (Corning: 10092CV) supplemented with 0.1% BSA, 1% B27 (v/v), ascorbic acid, glutamine, MTG, HGF (20ng/ml), and Epidermal growth factor (EGF, 50ng/ml, R&D: 236-EG) for 4 days to initiate cholangiocyte differentiation. To induce CFTR expression, the medium was replaced with DMEM/F12 medium containing 0.1% BSA, 1% B27 (v/v), ascorbic acid, glutamine, MTG, and Retinoic Acid (2μM, Sigma: R2625) for an additional 6 days. For maturation and the promotion of primary cilia formation, cells were further cultured for 12 days in DMEM/F12 medium with 0.1% BSA, 1% B27 (v/v), ascorbic acid, glutamine, MTG, Noggin (50ng/ml, R&D: 3344-NG), RI (5μM), and Forskolin (FSK, 5μM, TOCRIS: 1099). Media were refreshed every two days throughout the differentiation protocol. All culture were maintained under ambient oxygen conditions.

### Single-cell RNA-seq data processing and analysis

Single-cell RNA-sq (scRNA-seq) was performed on both XAV and CHIR-treated mature hPSC-hep 3D aggregate at day 53 following 18 days of maturation periods from hepatic progenitor cells. ScRNA-seq data associated with both XAV and CHIR treated mature hPSC-hep 3D aggregates are deposited in GEO: GSE299658. Sample processing was conducted according to the 10X Genomics Single Cell 3’ v2 Reagent User Guide and as previously described in the literature^50^. Briefly, sequencing library generation, data filtering to exclude cells with a high mitochondria ratio (>4 SD above means) and low library size (>1500), and normalization using the scran R package with default settings was performed^67^. Cell clustering using Seurat with standard procedures and differential expression analysis, as previously described^50^, identified 3 primary clusters in XAV-treated cells, and 2 primary clusters in CHIR-treated cells, respectively. Differential gene expression lists for each cluster are provided in Supplementary Data 1-5. To identify the cell types that were generated from the hPSC-derived hepatocytes, we performed Harmony integration^68^ with the human liver cell map described by MacParland et al. (GEO: GSE115469)^50^. Briefly, the raw count matrices for 10X single-cell experiments of both human primary liver cells and the hPSC-derived hepatocyte promoting the maturation with Wnt antagonist and agonists were merged and processed by our standard pipeline into one Seurat Single Cell Experiment object. PCA was first performed with the raw data to be used for integration. The hPSC-derived hepatocytes were integrated with human primary liver cells (as 2 groups: xenograft and primary) using the R package ‘‘harmony’’ (version 0.0.0.9000) using the following parameters: theta = 4, plot convergence = T, nclust = 15, max.iter.cluster = 100. The resulting clusters were determined by Seurat FindClusters with reduction (type = ‘‘harmony,’’ dims.use = PC1:PC30, and resolution = 0.2). The clusters were visualized by plotting tSNE reduction values on a Cartesian graph and UMAP.

To assess the similarity between hPSC-derived hepatocytes and various primary hepatocyte cell types, we performed Pearson correlation analysis using liver cell type signatures. This gene list represents the genes whose expression best distinguished the human liver cell types from each other. From this gene list, we focused our analysis on the differentially expressed genes of primary human hepatocytes (clusters 1,2, 5, 7,17, and 18) and hPSC-derived hepatocytes (clusters 10,15 and 19), from which we generated a merged and differentially expressed gene was determined and listed in Supplementary Data 7 by in Figure 5d and e.

Differential gene expression was calculated using the Wilcoxon Test in R with hPSC-derived hepatocyte clusters (cluster 10,15 and 19) compared to the rest of the population and the results were converted to ranking scores (rnk files) specific to each cluster. Rnk files were then used for pathway enrichment analysis by Gene Set Enrichment Analysis (GSEA)^69^ software from the Broad Institute with a custom pathway database Human_GOBP_AllPathways_no_GO_iea_August_01_2018_symbol.gmt (www.baderlab.org/GeneSets). Significant GSEA pathways were visualized using the Enrichment Map App in Cytoscape^70^ (Figure 5e, Supplemental Figure 5c, 6).

### Flow cytometry

Differentiated cells were dissociated into single-cell suspensions. Dead cells were excluded from analysis, and gating was determined using isotype controls (ALB and AFP) or unstained controls. For cell surface marker analysis, cells were stained in PBS supplemented with 10% FCS. For intracellular protein detection, cells were fixed in 4% paraformaldehyde (Electron Microscopy Science, Hatfield, PA, USA) in PBS, then permeabilized with 90% ice-cold methanol for 10 minutes for markers including ALB, AFP, and Ki67. After permeabilization, cells were incubated with secondary antibodies for 30 minutes at room temperature. Stained cells were analyzed using an LSRFortessa (BD), and data were processed using FlowJo software version 10 (Tree Star). FACS antibodies and their dilution ratios are provided in Supplementary Tables 1,2.

### ELISA

ALB levels were quantified using a human albumin ELISA kit (Bethyl Laboratories), following the manufacturer’s instructions. Briefly, 96-well plates were pre-coated with anti ALB antibody. After blocking, samples and standards were added to the wells and incubated for 1 hour at room temperature. Following incubation, HRP-conjugated secondary antibody was applied and incubated for an additional hour. Detection was carried using TMB substrate for 15 minutes. The reaction was stopped, and absorbance was measured using a microplate reader. Albumin concentrations were calculated based on a standard curve.

### CYP3A4 functional assay

CYP3A4 enzymatic activity in hPSC(H9 and WT01)-derived mature hepatocyte was assessed using the p450-Glo CYP3A4 Assay Kit (Promega), according to the manufacturer’s instructions. Briefly, cells were incubated with the luciferin-based substrates at 37°C to allow enzymatic conversion. After incubation, the reaction mixture was transferred to a 96-well half-area plate. Detection reagent was then added, and the plate was incubated at room temperature to stabilize the luminescent signal. Luminescence was measured using a microplate reader. Genomic DNA was extracted using DNA isolation kit (Qiagen), according to the manufacturer’s protocol. Relative luminescence units (RLU) obtained from CYP assays were normalized to DNA content to account for variations in cell number.

### Immunostaining

For staining of cells in tissue culture plates, cells were fixed with 4% paraformaldehyde for 15 min and permeabilized with either 0.2% Triton X-100 or cold 100% methanol (for Ki67 and Ku80) prior to blocking. Cells were washed three times with PBS for 5 minutes at room temperature before and after each staining step. Antibodies were diluted in DPBS supplemented with 0.1% BSA and 0.1% Triton X-100. For immunostaining of paraffin-embedded sections, sections were first dewaxed in xylene, rehydrated, and placed in Tris-EGTA buffer (10 mM Tris, 0.5 mM EGTA, pH 9.0) for heat-induced epitope retrieval for 20 minutes before blocking. Antibodies were diluted in DPBS containing 0.3% BSA and 0.3% Triton X-100.

Cells were stained with primary antibodies in staining buffer overnight at 4°C, followed by incubation with secondary antibodies for 1 hour at room temperature, and counterstained with DAPI. Stained cells were examined using an EVOS Microscope (Thermo Fisher Scientific) or a NIKON A1 Resonant Confocal Microscope, with images captured via Nikon Elements software. Details of antibodies used are provided in Supplementary Tables 1,2.

### Quantitative real-time PCR

Total RNA was isolated using the RNAqueous Micro Kit (Invitrogen). Between 500 ng and 1 μg of RNA was reverse transcribed into cDNA using the iSCRIPT Reverse Transcription Supermix (Bio-Rad). Quantitative PCR was performed on a C1000 Touch Thermal Cycler (Bio-Rad) using the SsoAdvanced Universal SYBR Green Supermix (Bio-Rad). Gene expression levels were normalized to the housekeeping gene TATA box binding protein (TBP). Oligonucleotide sequences are provided in Supplementary Table XX, and control RNA samples from adult liver (AL), fetal liver (FL), and pancreas (PANC) are listed in Supplementary Table 3,4.

### Transmission Electron Microscope (TEM)

Samples were fixed in 2.5% glutaraldehyde in PBS, rinsed and post-fixed in 1% osmium tetroxide (Electron Microscopy Sciences) for 1 hour. Following an additional rinse with 0.1M Sorenson’s phosphate buffer, the tissue was dehydrated through an ascending ethanol series, infiltrated with and embedded in modified Spurr’s resin. The region of interest was first identified using thick sectioning, and then ultrathin sections (90-100nm) were obtained with a Leica UC6 ultramicrotome (Leica). The sections were subsequently stained with Uranyless and lead citrate, then imaged using a Hitachi HT7700 transmission electron microscope (Hitachi). Analysis was performed on samples derived from three independent experiments.

### Seahorse assay

For the Seahorse XF Mito Stress Test, twelve hPSC(H9)-derived magure 3D hepatic aggregates or primary hepatocyte aggregates were plated onto an XFe96 cell culture microplate pre-coated with 2.5% Matrigel. Each organoid contained 2000 cells. Forty-five minutes prior to the assay, cells were incubated in Seahorse XF DMEM assay medium (Agilent) supplemented with 10mM Glucose, 1mM pyruvate, and 2mM glutamine at 37 °C. The assay cartridge was preloaded with XF Cell Mito Stress Test compounds (3μM oligomycin, 4μM FCCP, 1μM rotenone/1μM antimycin A) 30 min before the assay. Following calibration, the XF Cell Culture Microplate was inserted into the Seahorse XFe Analyzer, and the XF Cell Mito Stress Test was conducted. After measuring the oxygen consumption rate (OCR), cells were lysed for DNA content normalization using the CyQUANT Cell Proliferation Assay kit, following the manufacturer’s instructions (Thermo Fisher Scientific). Data were analyzed using Wave software (Agilent).

### Primary human hepatocytes

Non-cryopreserved human primary hepatocytes (HepaSH)^58^ were transported from CIEM (Japan) to Toronto. Upon arrival, cells were washed, and dead cells were removed with Percoll following the manufacturer’s instructions. Cell viability was 93.32 ± 1.15% (mean ± SEM, *n* = 10). For Seahorse assays, 2,000 hepatocytes per well were seeded into low-attachment 96-well U-bottom plates to form aggregates, and 12 aggregates were plated per well of seahorse XFe96 cell culture microplate two days prior to the assay. For CYP3A4 activity assays, 1 million cells per well of 6 well EZSPHERE SP Microplate Plate (IWAKI, 4810-900SP) were seeded to make aggregates for two days and cells were transferred to 6-cm dishes. Assays were conducted at the indicated time points in the figure. For ELISA and 3D printing, cell were aggregated in the EZSPHERE SP Microplate Plate as CYP3A4 assay and maintained in HCM media (LONZA, CC3198).

### 3D Bioprinting

HB, XAV-treated and CHIR treates hepatic aggregates generated from hPSC (H9) were harvested, pelleted, and resuspended at 2 X10^7^ cells per 100ul of medium. The cells suspension was combined with 1ml of LAMININK bioink (CELLINK: IKC20500001) at the ratio of 1ml bioink per 20 million cells,gently mixed, and loaded into a sterile cartridge for extrusion on a BIO X^TM^ printer (CELLINK; DNA Studio v4.1.3). Grid-like constructs (10mm X 10mm X1mm) were printed using the follow parameters: nozzle pressure 3–5 kPa, extrusion speed 3 mm s⁻¹, and 500 ms pre-flow. Immediately after printing, lattices were cross-linked in 100 mM CaCl₂ for 5 min at room temperature, rinsed with culture medium, and transferred to standard incubation conditions. This protocol yields up to eight constructs per millilitre of bioink, each containing ∼2.5 × 10⁶ viable cells; minor variation reflects residual bioink retained within the nozzle.

### *In vivo* animal experiment

For renal subcapsular implantation of hPSC (H9)-derived 3D hepatic aggregates, NSG mice (10-15 weeks old; *n* = 5; 3 males and 2 females) were anesthetized with isoflurane. Following a small flank incision, the left kidney was externalized, and a superficial incision was made in the caudal region of the kidney capsule using a 25G needle. A polyethylene catheter (PE50) preloaded with approximately 1 × 10⁷ cells (HBs, XAV-treated, CHIR-treated, or a 1:1 mixture of XAV and CHIR-treated aggregates) suspended in 50–100 µL of culture medium was inserted into the subcapsular space. Peripheral blood was collected at 2 and 4 weeks post-implantation to assess human albumin secretion by enzyme-linked immunosorbent assay (ELISA). Grafts were harvested at week 4 for histological and immunohistochemical analysis. For intrahepatic transplantation, liver injury was induced in TK-NOG mice (7–10 weeks old, male only) by intraperitoneal injection of ganciclovir (GCV; Sigma G2536) at a dose of 25 mg/kg, administered once, 7 days prior to cell transplantation. Mice that did not exhibit sufficient liver injury as determined by subthreshold elevations in plasma alanine aminotransferase (ALT) and alkaline phosphatase (ALP) levels, received an additional intraperitoneal dose of ganciclovir (10 mg/kg) to ensure sustained hepatic damage. Cell transplantation was then performed via splenic injection of 1.0 × 10⁶ cells suspended in 100 µL of sterile saline. Transplanted cell populations included hepatic progenitor cells (HBs) derived from hPSC (H9) (*n* = 10), day 18–24 XAV-treated hepatocyte-like cells (*n* = 15), and day 18–24 CHIR-treated hepatocyte-like cells (n = 15), all obtained from enzymatically dissociated monolayer cultures. Cryopreserved primary human hepatocytes (Lonza; HUCSD, Lot 181001A, 12-year-old female donor) served as positive controls (*n* = 4). Blood samples were collected at weeks 4 and 12 post-transplantation to quantify human albumin secretion. Liver tissues were harvested at weeks 12 and 16 for histological evaluation of engraftment and tissue integration. All animal procedures were conducted in accordance with protocols approved by the University Health Network Animal Resource Centre (UHN ARC). Mice were anesthetized using isoflurane and received Tramadol for postoperative analgesia. Postoperative care and housing complied with institutional animal care and use guidelines.

### Statics and reproducibility

Statistical methods and the numbers of biological replicates were detailed in the figure legends. All experiments were repeated at least three times independently. The standard error of the mean (SEM) was calculated using data from both biological and technical replicates. Statistical analyses were performed using GraphPad Prism 10.

## Acknowledgements

We gratefully acknowledge the members of the Ogawa, Keller, McGilvray, MacParland, and Bader laboratories for their valuable input on the experimental design and manuscript preparation. We also thank the University Health Network/SickKids Flow Cytometry Facility, the UHN Pathology Research Program, the Princess Margaret Genomics Centre, the Animal Resource Centre (ARC), and the Advanced Optical Microscopy Facility at the Princess Margaret Cancer Research Tower for their technical support. We extend our appreciation to Mr. Yongbo Bai (SCREEN Holdings Co., Ltd.) for his assistance with high-throughput imaging analysis during the culture of cryopreserved hepatoblasts in Elplasia 96-well plates (Corning 4442). We also thank Dr. Ravi Vellanki from the Wouters laboratory for technical support with Seahorse metabolic flux assays. We are especially grateful to Dr. Gordon Keller for his invaluable guidance throughout this work, including his review of this manuscript. Figure 6a, Supplementary Figure 8a, 10a, 11a were created with BioRender.com. This work was supported in part by the Medicine by Design Initiative at the University of Toronto, funded through the Canada First Research Excellence Fund (CFREF), the Canadian Institutes of Health Research (CIHR; Grant IDs: 1023968 and 1021677), and the Stem Cell Network through a Fuel Biotechnology Project Grant (Grant ID: 1025105).

## Author contributions

M.O. and S.O. contributed to the experimental design, data acquisition, analysis, interpretation, and manuscript writing. J.L. performed the scRNA-seq analysis. A.D., B.T., X.Z., K.M., K.T., C.C., M.H., and Y.H. conducted experiments. I.F. provided technical support. H.S., I.M., S.M., and G.B. offered experimental guidance. S.O. supervised the experiments, edited the manuscript, and approved the final version.

## Data availability

The data supporting this study are available within the Article and its Supplementary Information files. Raw scRNA-seq data generated in this study have been deposited in the GEO database under the accession code GE299658. Referenced human liver scRNA-seq data are available under the accession code GSE115469.

## Competing interestes

The authors declare no competing interests.

---

**Supplementary Figure 1.**
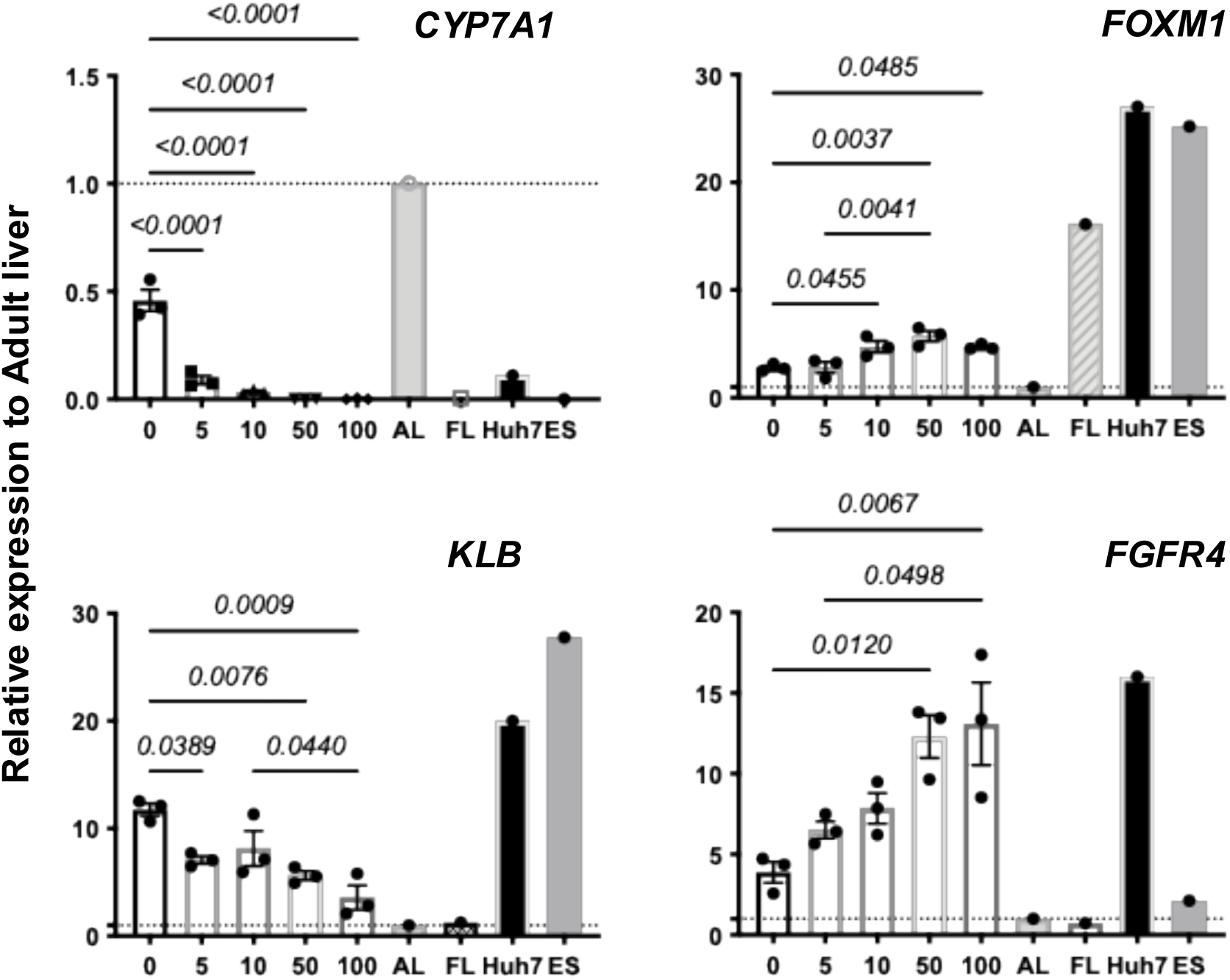
RT-qPCR analysis of gene expression in hepatic progenitors following dose-dependent FGF19 treatment. Hepatoblasts were treated with increasing concentrations of FGF19 (5,10,50, and 100 ng/ml). RT-qPCR data showing the expression of target genes. Expression levels were normalized to TBP and are presented relative to adult liver (set to 1). AL adult liver, FL fetal liver. One-way ANOVA. DATA are represented as mean ± SEM (*n* = 3).

**Supplementary Figure 2.**
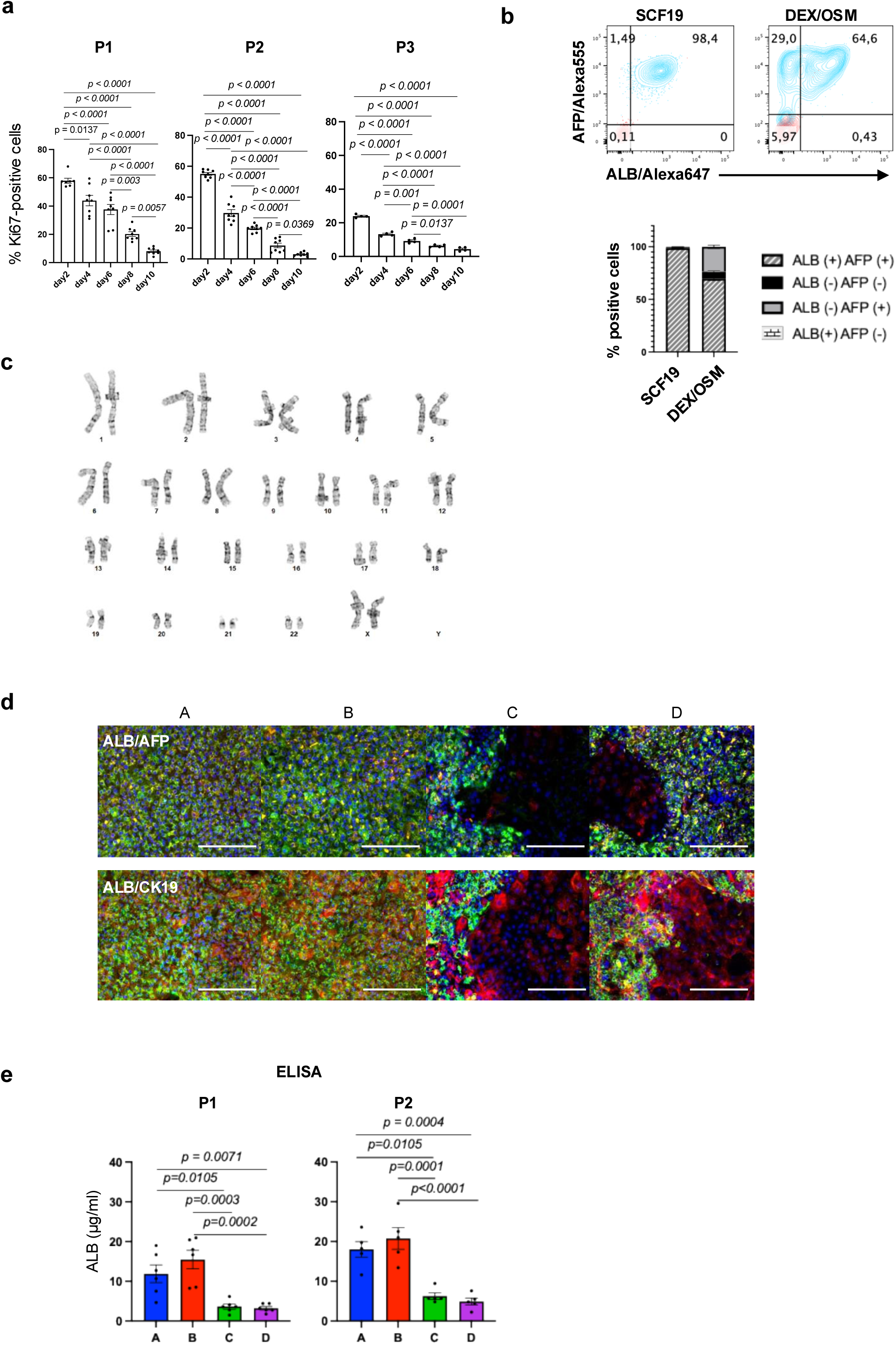
Serial expansion and cryopreservation of hPSC-derived hepatic progenitor cells in SCF19 medium. a. Quantification of Ki67^+^ cells at the indicated time points during passages cultured in SCF19 expansion cocktails. b. Flow cytometry analysis and its quantification of ALB^+^ and AFP^+^ cell populations following treatment with either SCF19 or DEX/OSM (*n* = 9). c. Karyotyping analysis showing a normal 46, XX karyotype in H9-derived hepatoblasts at passage 5. d. Immunostaining analysis showing the co-expression of ALB (green) with AFP (red), or ALB (green) with CK19 (red) in cells cultured in different expansion cocktails. Scale bar: 200µm. e. ELISA analysis showing ALB secretion from 3D suspension cultures grown in different expansion cocktails. 3D aggregates were generated from passage 1 (P1) and passage 2 (P2) monolayer cultures. One-way ANOVA. DATA are represented as mean ± SEM (*n* = 6 for P1, *n* = 5 for P2).

**Supplementary Figure 3.**
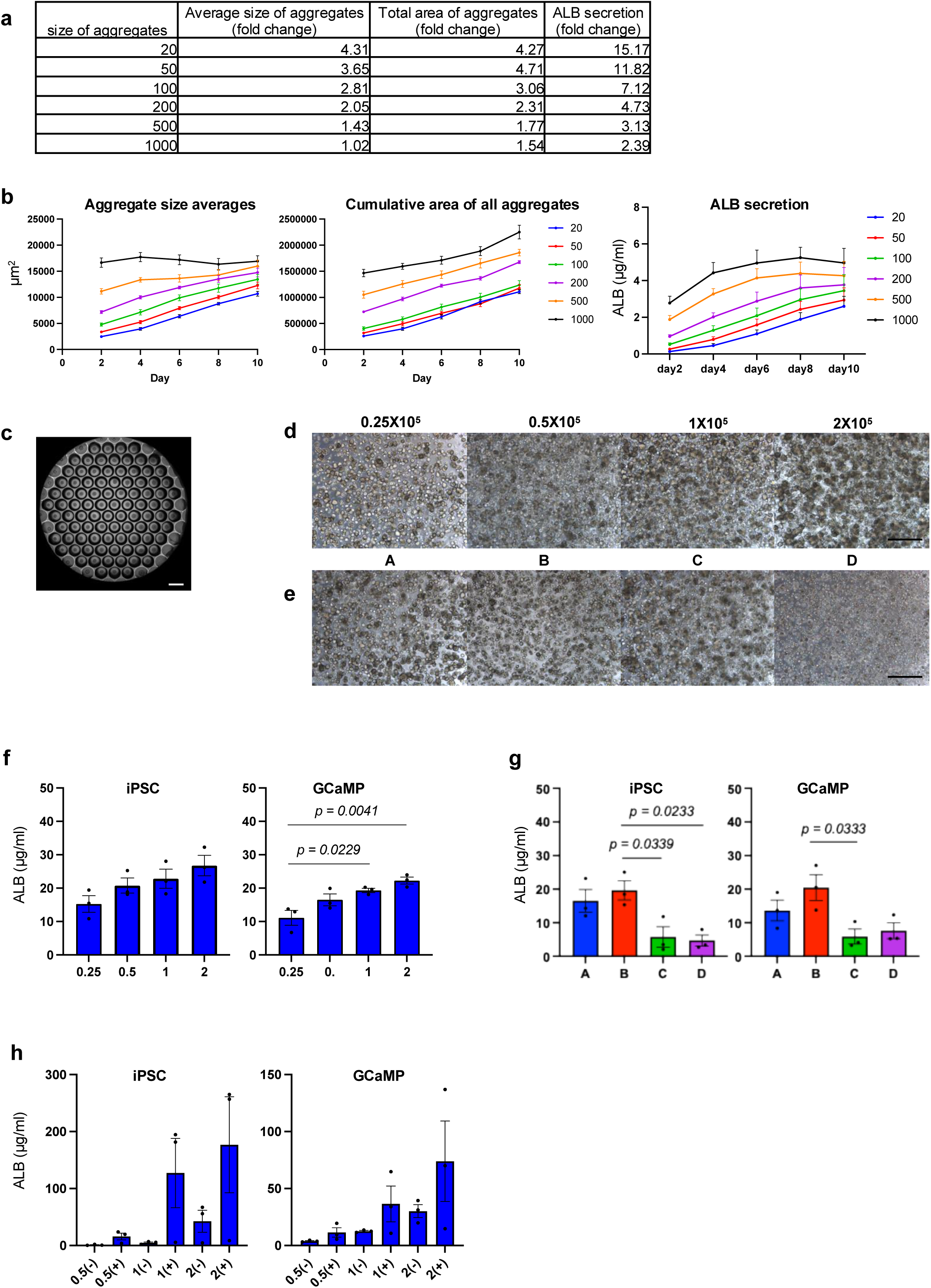
SCF19 medium facilitates scalable 3D aggregation of hPSC-derived hepatic progenitors. f. Table showing the fold change quantification of the indicated parameters in H9-derived hepatoblasts at different initial cell densities. g. Quantification of aggregate size averages, cumulative area of all aggregates, and ALB secretion in directly passaged H9-derived hepatoblasts aggregate cultures with different initial cell numbers over a 10-day period. h. Microscopic analysis of H9-derived 3D hepatoblasts organoids directly passaged in microwells. Scale bar: 500µm. i. Microscopic analysis showing H9-derived 3D hepatoblasts organoids cultured in Matrigel at different initial cell densities over a 10-day period. Scale bar: 500µm. j. Microscopic analysis showing H9-derived 3D hepatoblasts organoids cultured in Matrigel with an initial cell density of 0.25X10^5^, in the presence of different expansion cocktails over a 10-day period. Scale bar: 500µm. k. ALB secretion in iPSC and GCaMP-derived hepatoblasts in Matrigel culture at different initial cell densities. One-way ANOVA. DATA are represented as mean ± SEM (*n* = 3). l. ALB secretion in iPSC and GCaMP-derived 3D hepatoblasts cultured in Matrigel with different expansion cocktails. One-way ANOVA. DATA are represented as mean ± SEM (*n* = 3). m. ALB secretion in iPSC and GCaMP-derived 3D hepatoblasts cultured in suspension with or without Matrigel at different cell densities (*n* = 3).

**Supplementary Figure 4.**
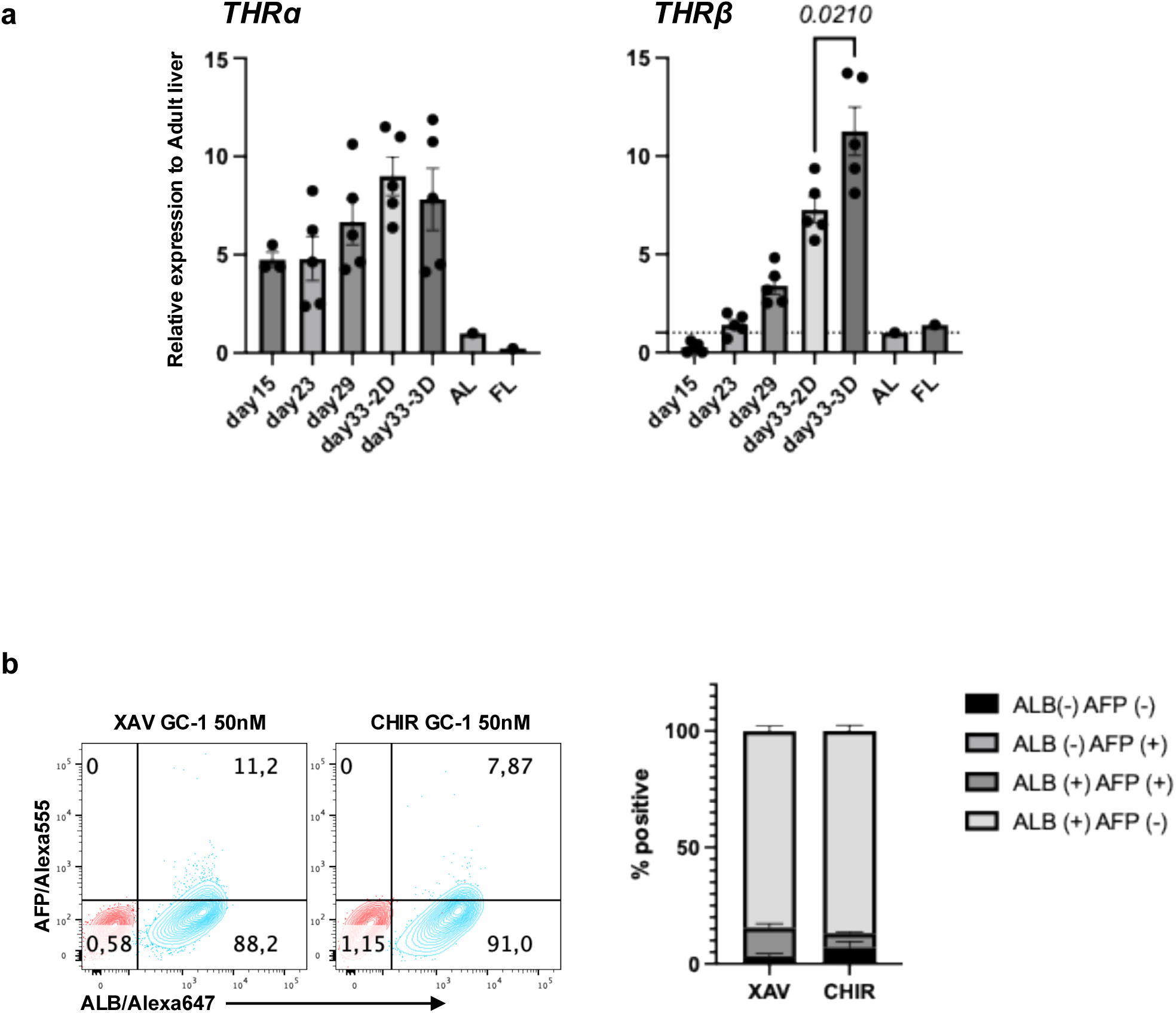
THRβ-selective agonist GC-1promotes maturation in 3D hepatic aggregates. a. RT-qPCR analysis of the expression of the indicated gene at different stage of differentiation. AL adult liver, FL fetal liver. One-way ANOVA. DATA are represented as mean ± SEM (*n* = 3-5). b. Flow cytometry analysis showing the population of ALB^+^ and AFP^+^ cells in GC-1 with XAV939 or CHIR99201-treated hepatic aggregates (left). Quantification of positive cells in each quadrant (right). DATA are represented as mean ± SEM (*n* = 5).

**Supplementary Figure 5.**
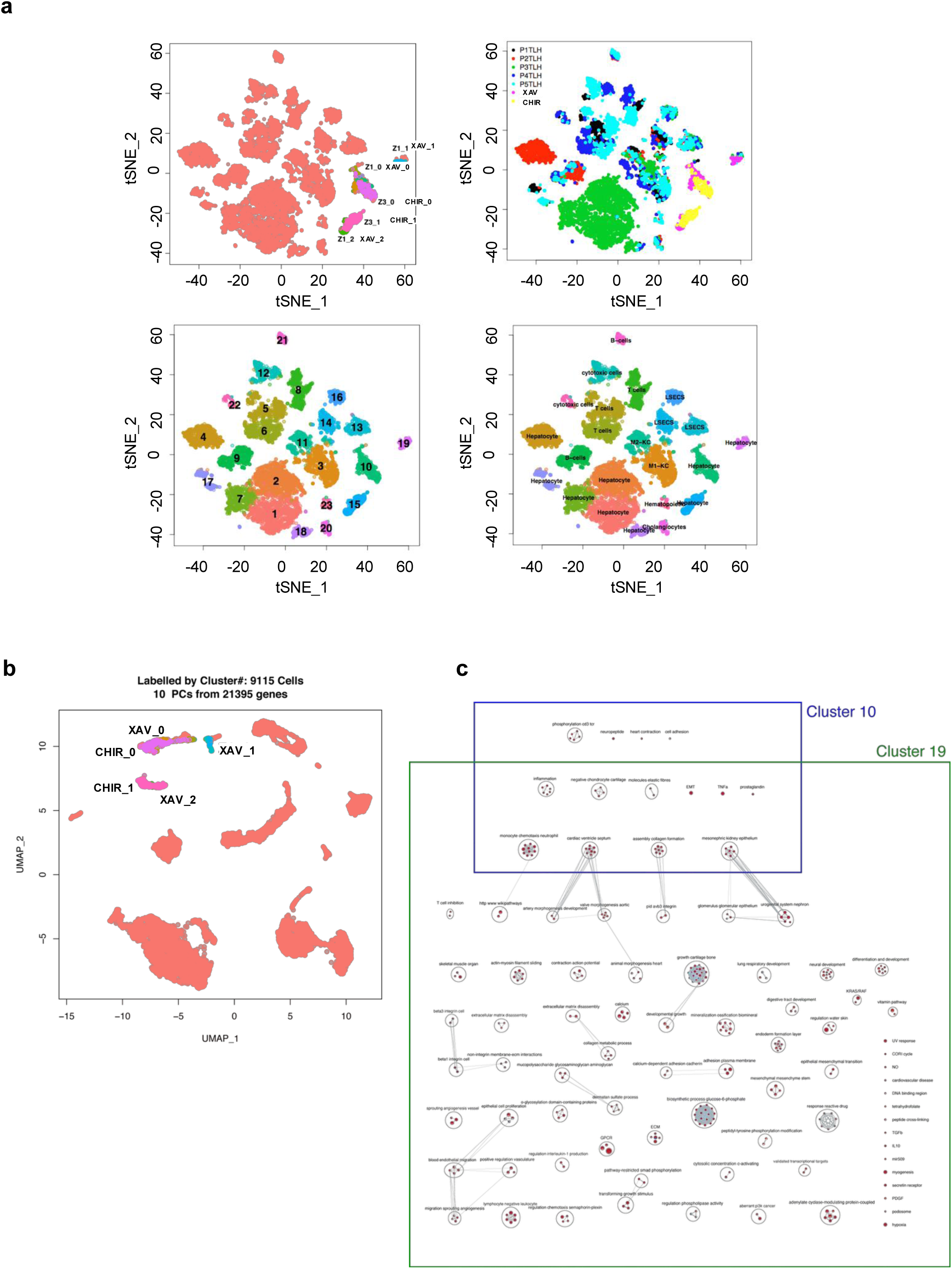
Comparative transcriptomic analysis of H9-derived hepatic aggregates and adult human liver. a. tSNE projection of integrated single-cell transcriptomic data from five adult human liver tissue samples (total liver homogenates) and H9-derived hepatocytes treated with either XAV or CHIR. A total of 9,115 cells were analyzed, encompassing 7,562 genes, and computationally resolved into 23 transcriptionally distinct clusters. **Top left:** Visualization of clusters enriched in human liver-derived cells (pink dots). **Top right:** Overlay of hPSC-derived hepatocyte populations: XAV-treated (pink dots) and CHIR-treated (yellow dots) to assess integration and cellular identity. **Bottom left:** Cluster assignment map displaying numerical labels for all 23 clusters. **Bottom right:** Cell type annotation based on canonical marker gene expression, allowing inference of hepatocyte, cholangiocyte, endothelial, and non-parenchymal cell lineages across clusters. b. UMAP projection of human adult liver cells and 3D H9-derived hepatic aggregates labeled by cell cluster. c. Pathway enrichment analysis comparing H9-derived hepatocyte clusters (Clusters 10 and 19) to adult primary hepatocyte clusters (Clusters 1, 2, 4, 7, 17, and 18), highlighting signaling pathways significantly upregulated in hPSC-derived populations.

**Supplementary Figure 6.**
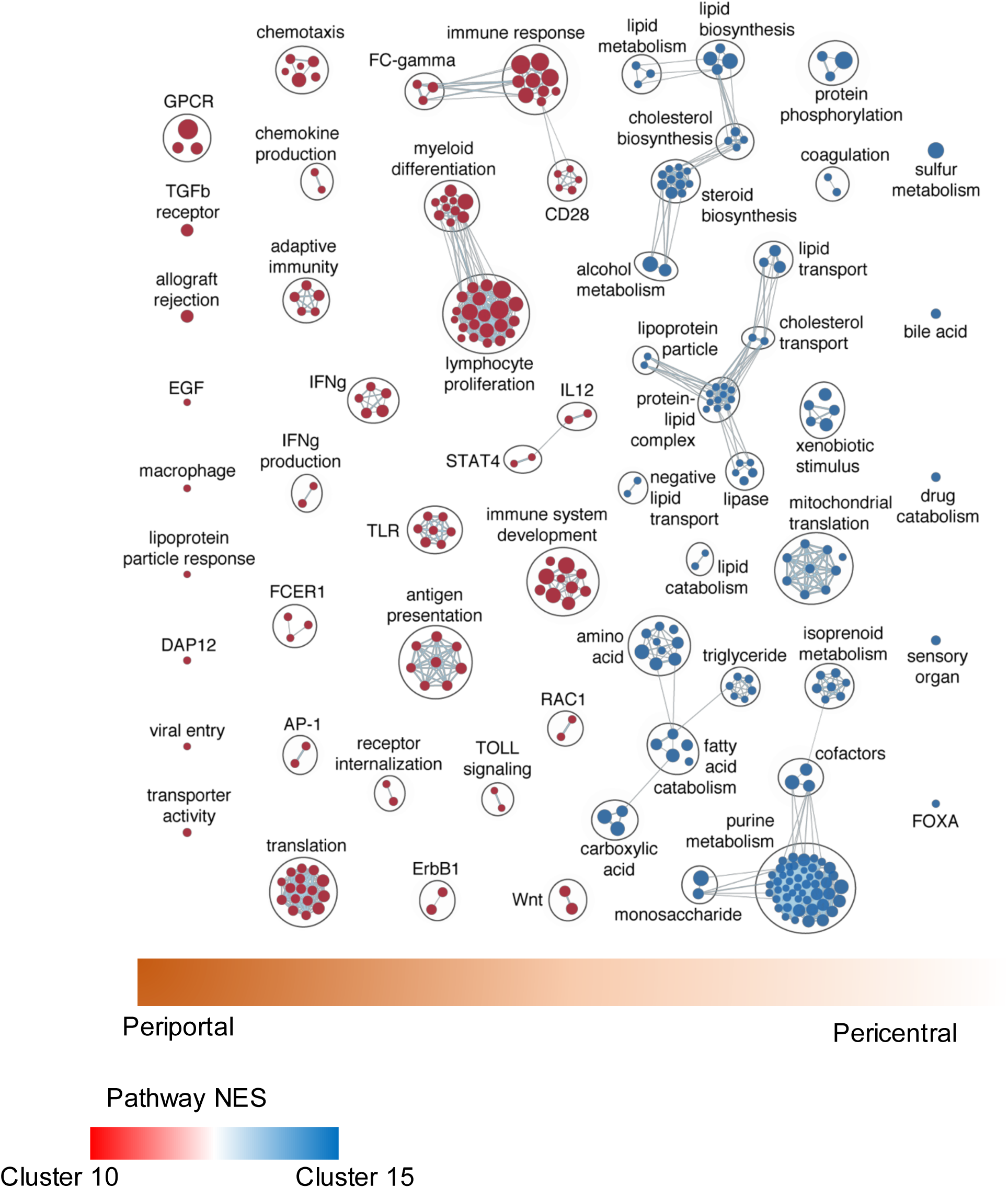
Signaling Pathways Upregulated in Zonation-Associated Clusters 10 and 15 Identified by Pathway Analysis.

**Supplementary Figure 7.**
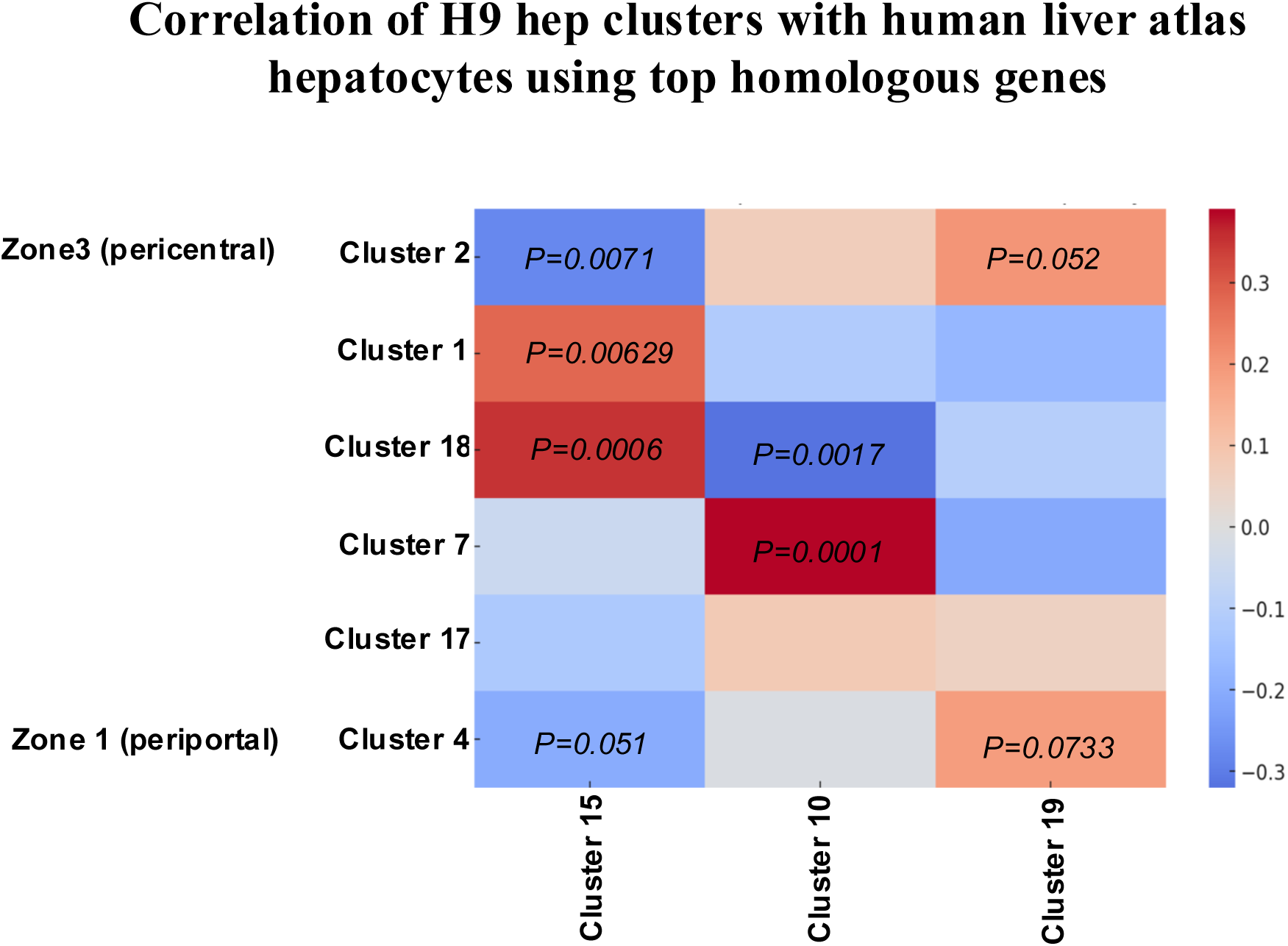
Transcriptional correlation between adult human liver and H9-derived hepatic aggregates. Heatmap illustrating the transcriptional similarity across clusters derived from adult human liver and H9-derived hepatic aggregates. Correlation coefficients were calculated based on the expression of top variable genes, highlighting zones of convergence and divergence between *in vitro*-derived hepatocyte populations and *in vivo* hepatic tissue. The color gradient represents the degree of correlation, with higher values indicating stronger similarity between clusters. Clustering is based on top homologous genes.

**Supplementary Figure 8.**
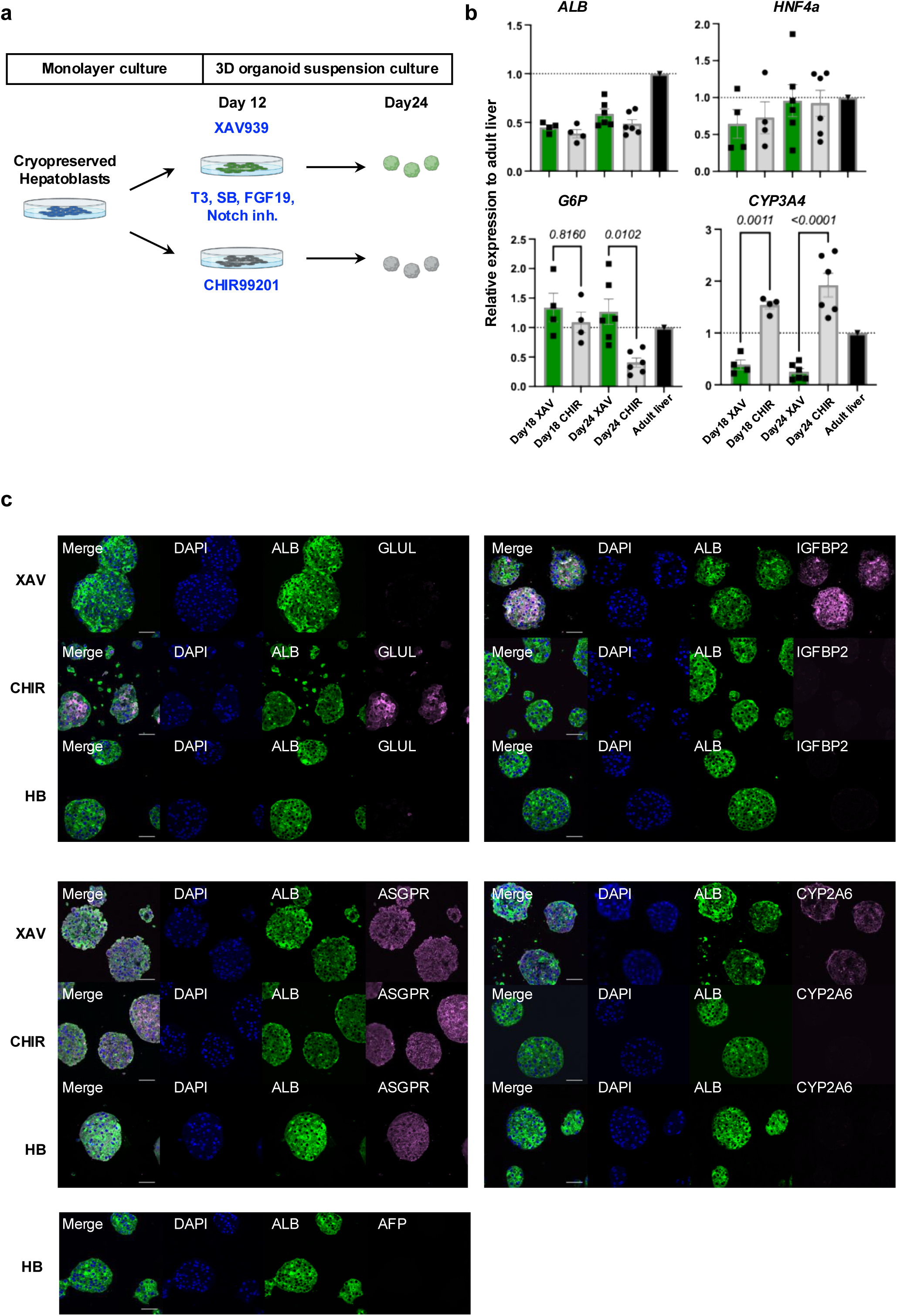
Functional differentiation of cryopreserved H9-derived hepatic progenitors into mature hepatocytes. a. Schematic differentiation protocol of hPSC-derived hepatocytes from cryopreserved hepatoblasts. b. RT-qPCR analysis of gene expression in hepatic cells treated with XAV939 or CHIR99021 at different time point of differentiation. AL adult liver. One-way ANOVA. DATA are represented as mean ± SEM (*n* = 4-6). c. Immunostaining analysis in hepatic aggerates treated with XAV939 or CHIR99021 showing the indicated proteins. Scale bar: 50µm.

**Supplementary Figure 9.**
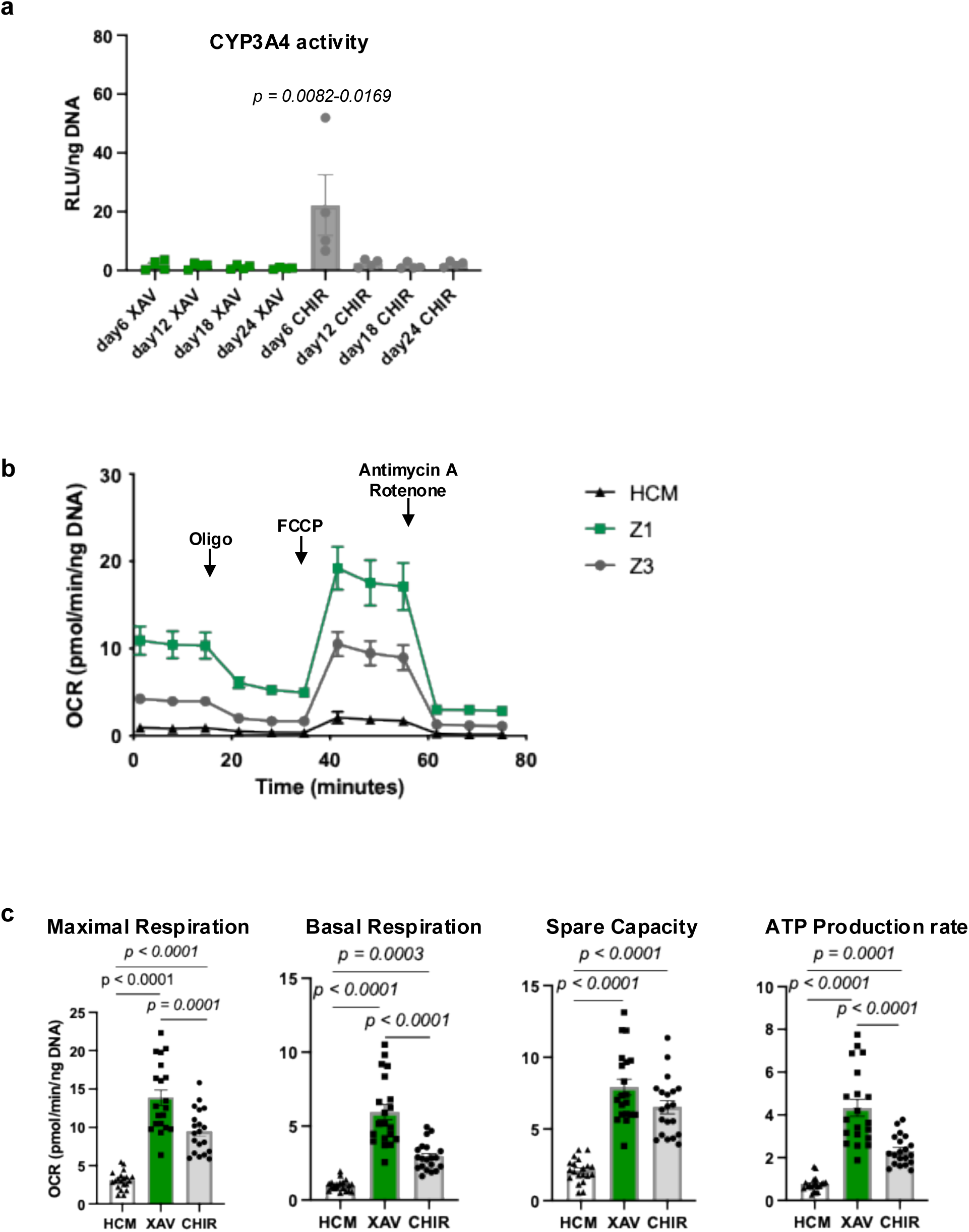
Functional characterization of primary human hepatocytes (HepaSH) via metabolic profiling a. CYP3A4 activity in different treated PH-derived hepatic aggregates at day6, 12, 18 and 24. Day6 CHIR showed significantly higher values compared to all other groups One-way ANOVA. DATA are represented as mean ± SEM (n = 5). Significant differences are indicated in the figure. b. Representative kinetics of the oxygen consumption rate (OCR) measured in primary hepatocyte aggregates cultured in indicated conditions. c. Comparison of each parameter in primary hepatocyte aggregates. One-way ANOVA. DATA are represented as mean ± SEM from 4 technical replicates of 5 biologically independent experiments.

**Supplementary Figure 10.**
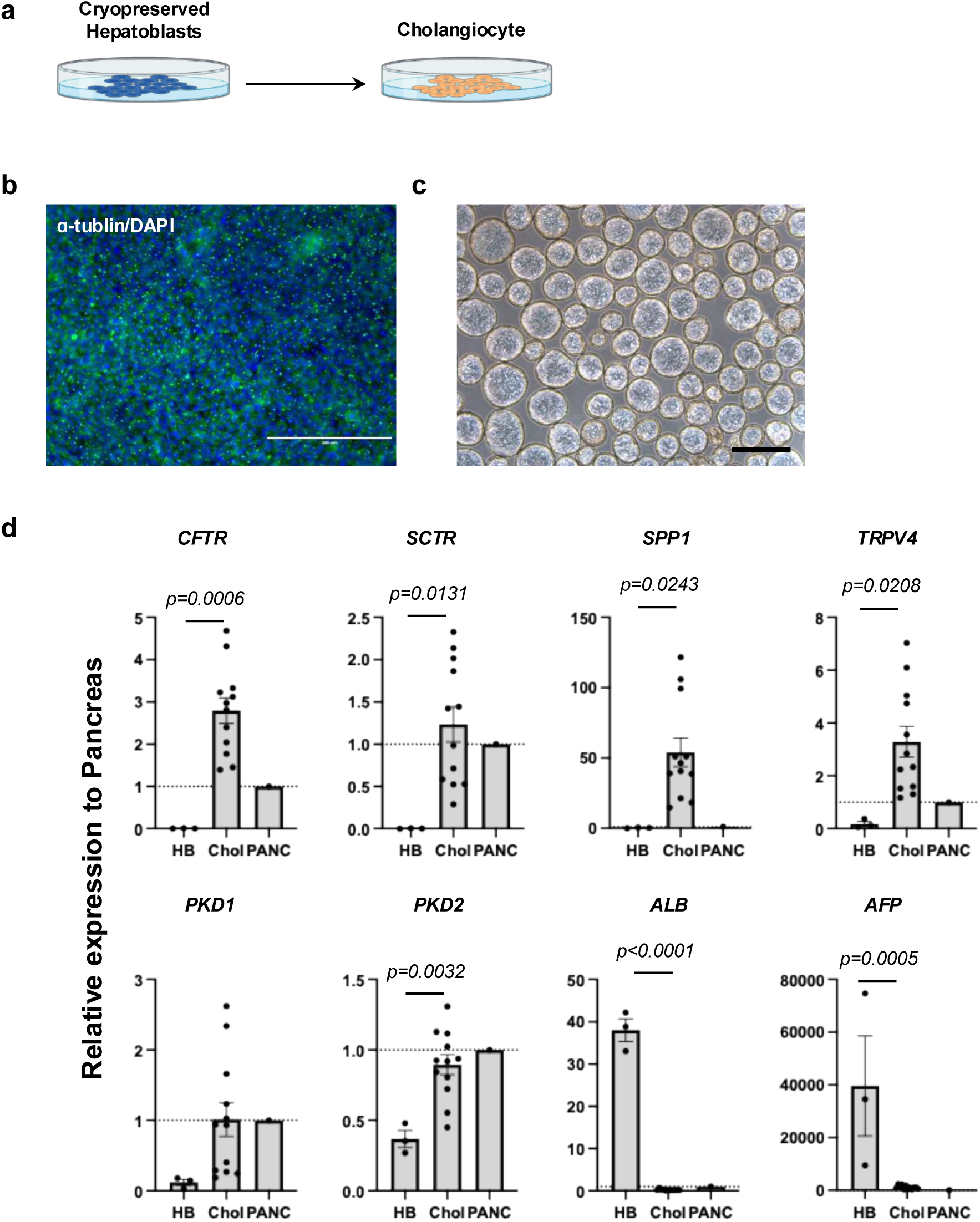
Differentiation of cryopreserved hPSC-derived hepatoblasts into functional cholangiocytes. a. Schematic differentiation of hPSC-derived cholangiocytes from cryopreserved hepatoblasts. b. Immunostaining analysis showing the proportion of cilia (green), and DAPI (blue) in cryopreserved iPSC HB-derived cholangiocytes. Scale bar: 200µm. c. Microscopic analysis of cryopreserved GCaMP HB-derived cholangiocyte cysts. Scale bar: 500µm. d. RT-qPCR analysis of the expression of the indicated gene cryopreserved HB-derived cholangiocyte. Expression levels were normalized to TBP and are presented relative to pancreas (set to 1). PANC pancreas. DATA are represented as mean ± SEM (HB; *n* = 4, Chol; *n* = 12).

**Supplementary Figure 11.**
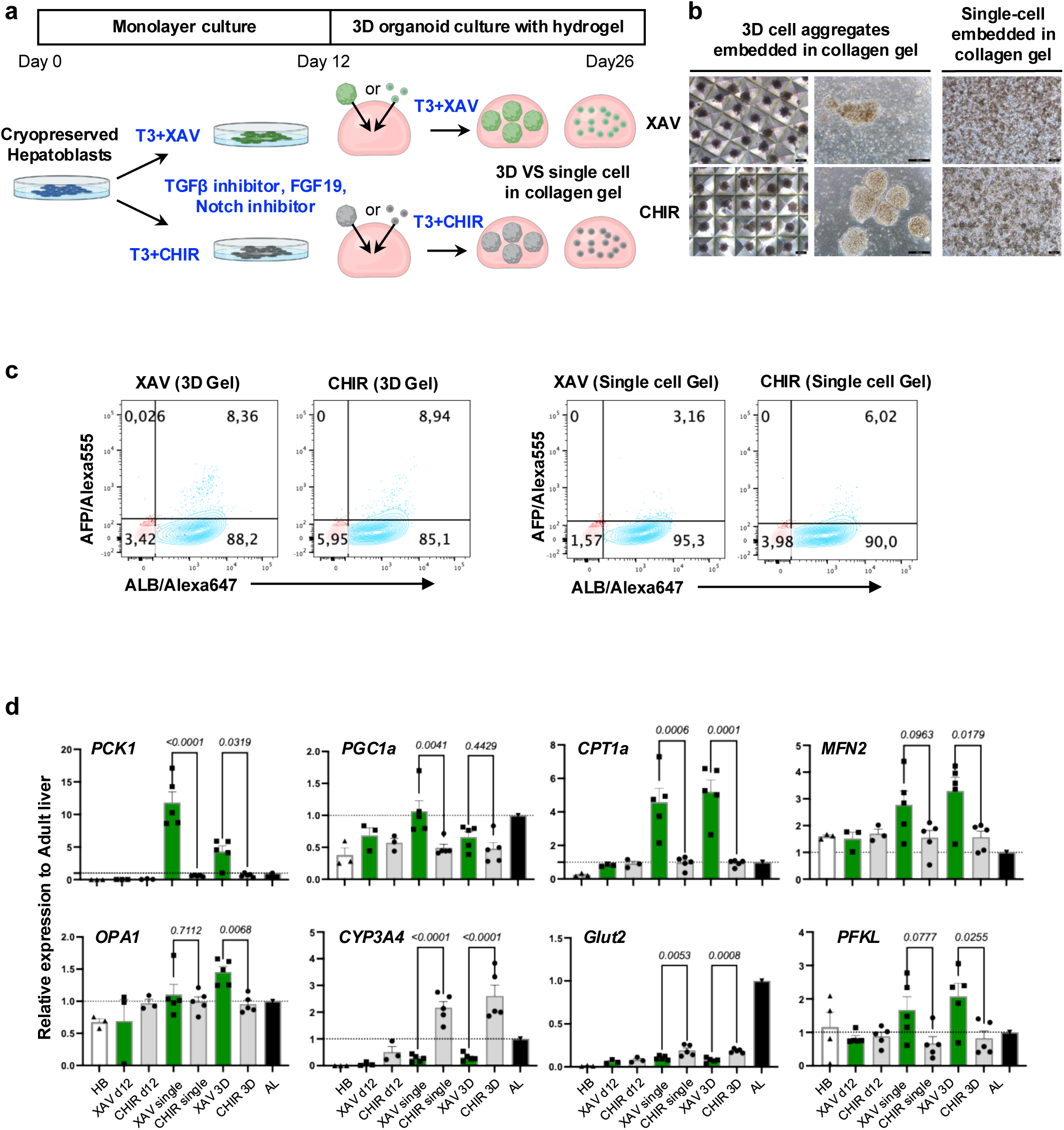
Wnt signaling modulation differentially regulates functional maturation and mitochondrial gene expression in hepatic aggregates derived from cryopreserved H9-derived hepatoblasts. a. Schematic differentiation protocol of H9-derived hepatocytes from cryopreserved hepatoblasts. b. Microscopic analysis showing hepatic aggregates in aggrewells (left), embedded in collagen (middle), and single cells in collagen (right) scale bars: 200µm. c. Flow cytometry analysis showing the population of ALB^+^ and AFP^+^ cells with XAV939 or CHIR99021-treated hepatic cells. d. RT-qPCR analysis of gene expression in hepatic cells treated with XAV939 or CHIR99021 across various culture formats. AL adult liver. One-way ANOVA. DATA are represented as mean ± SEM (*n* = 3-5).

**Supplementary Figure 12.**
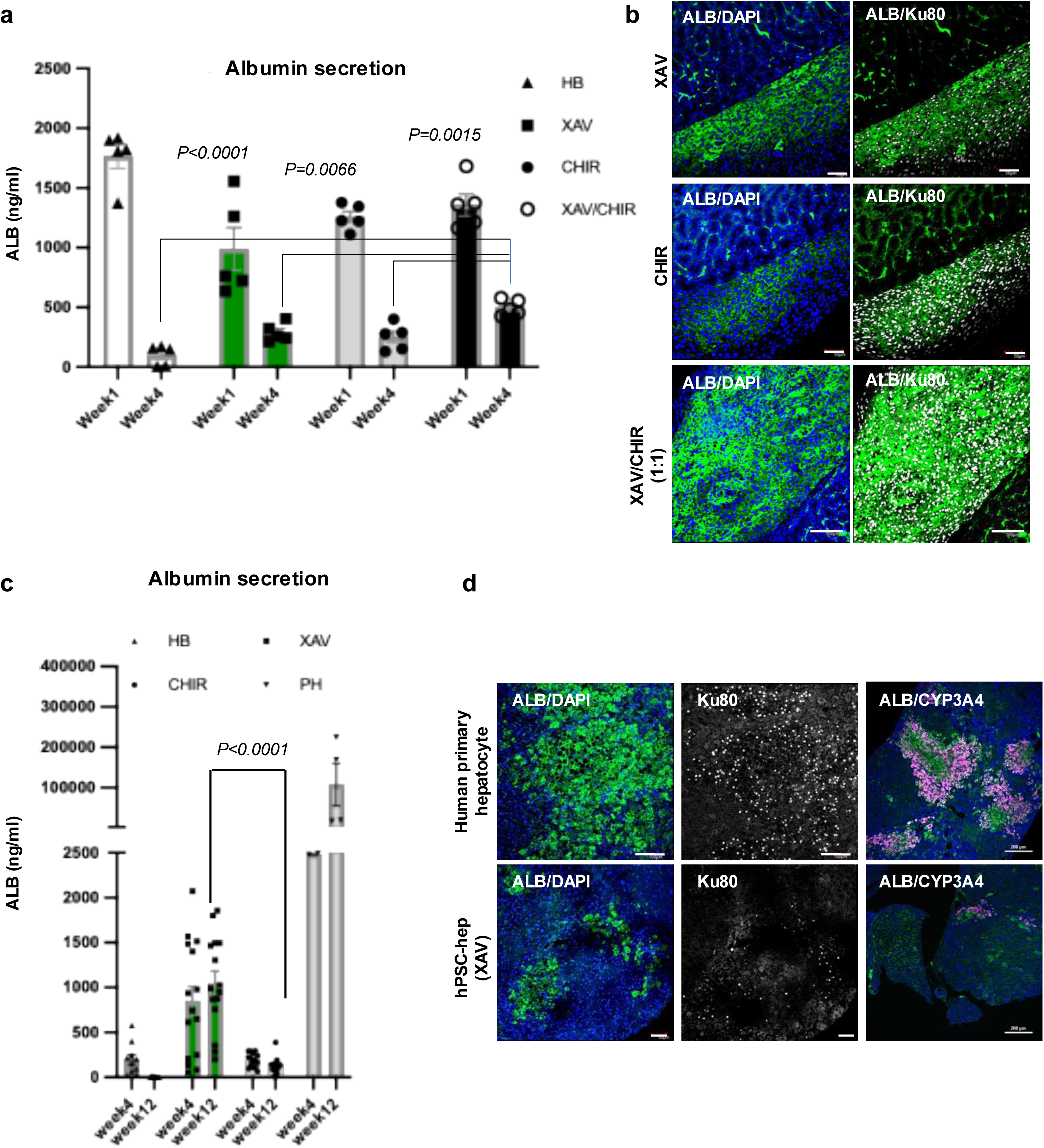
*In Vivo* functional assessment of H9-derived hepatocytes following transplantation. a. Human ALB secretion in the kidneys of NSG mice 4 weeks after transplantation with hepatic cells treated with XAV939 or CHIR99021. ALB secretion were compared among groups at week 4 using One-way ANOVA. Statistical significance is indicated in the figure. DATA are represented as mean ± SEM (n = 5). b. Confocal image of histological section in the kidney of NSG mouse after 4weeks of transplantation expressing human ALB and Ku80. Scale bar: 50 µm (top and middle), 400 µm (bottom). c. Human ALB secretion in TK-NOG mice post-transplantation of hepatic cells treated with XAV939 or CHIR99021. One-way ANOVA. DATA are represented as mean ± SEM (n = 15 in XAV and CHIR, *n* = 10 in HB). d. Confocal image of histological section in the liver of TK-NOG mouse after 16 weeks of transplantation of human primary hepatocytes (top row) and H9-derived XAV-treated hepatic cells (bottom row), expressing human ALB (green) and Ku80 (white) Scale bar: 100 µm (top), 50µm (bottom). ALB (green) and CYP3A4 (magenta). Scale bar: 200µm.

**Supplementary Figure 13.**
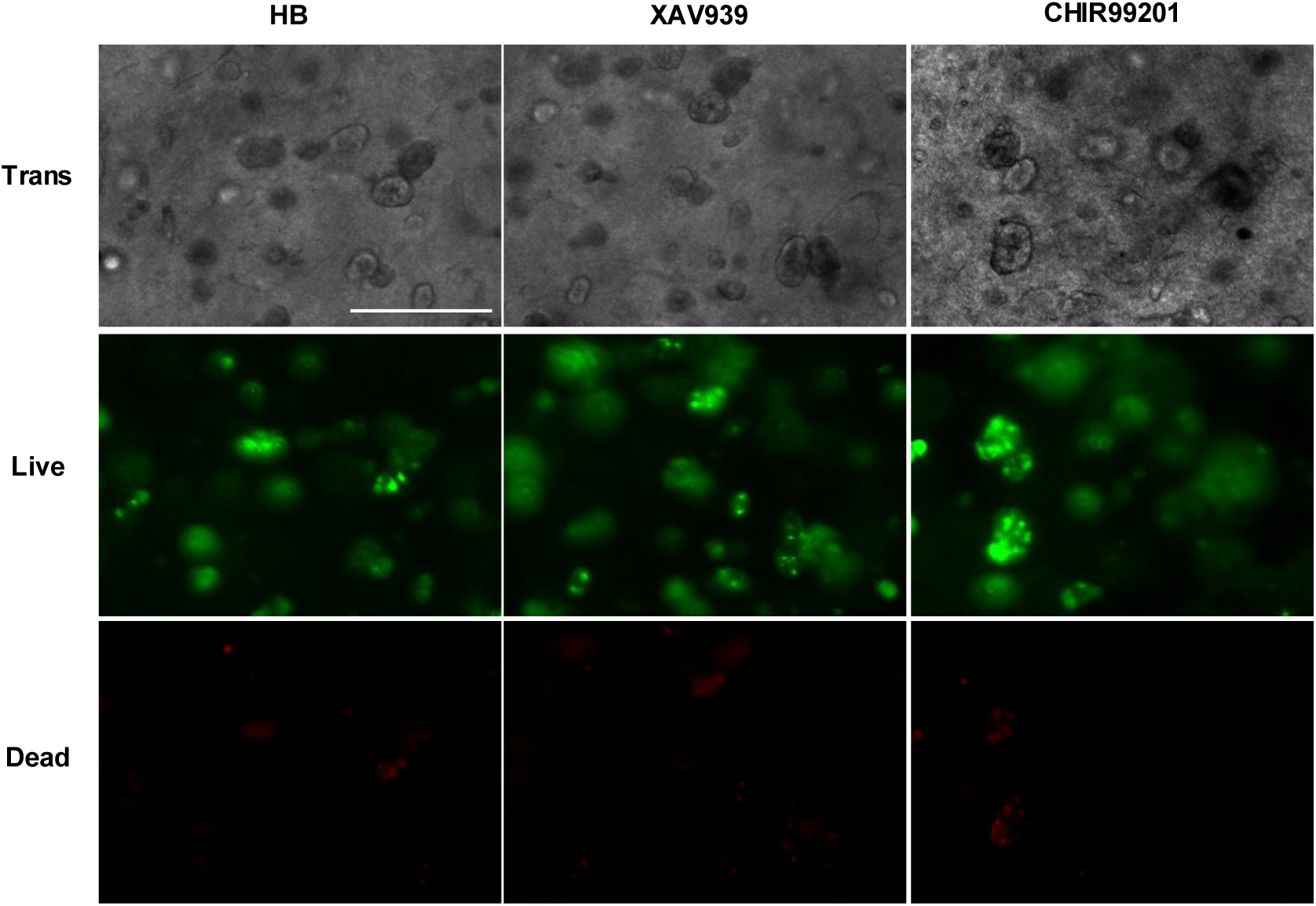
Microscopic visualization of 3D bioprinted hepatic constructs with corresponding live/dead viability staining. The top row displays phase-contrast, The middle row presents live staining (green fluorescence) of viable cells, and bottom row shows dead staining (red fluorescence). Scale bar: 200µm.

**Supplementary Figure 14.**
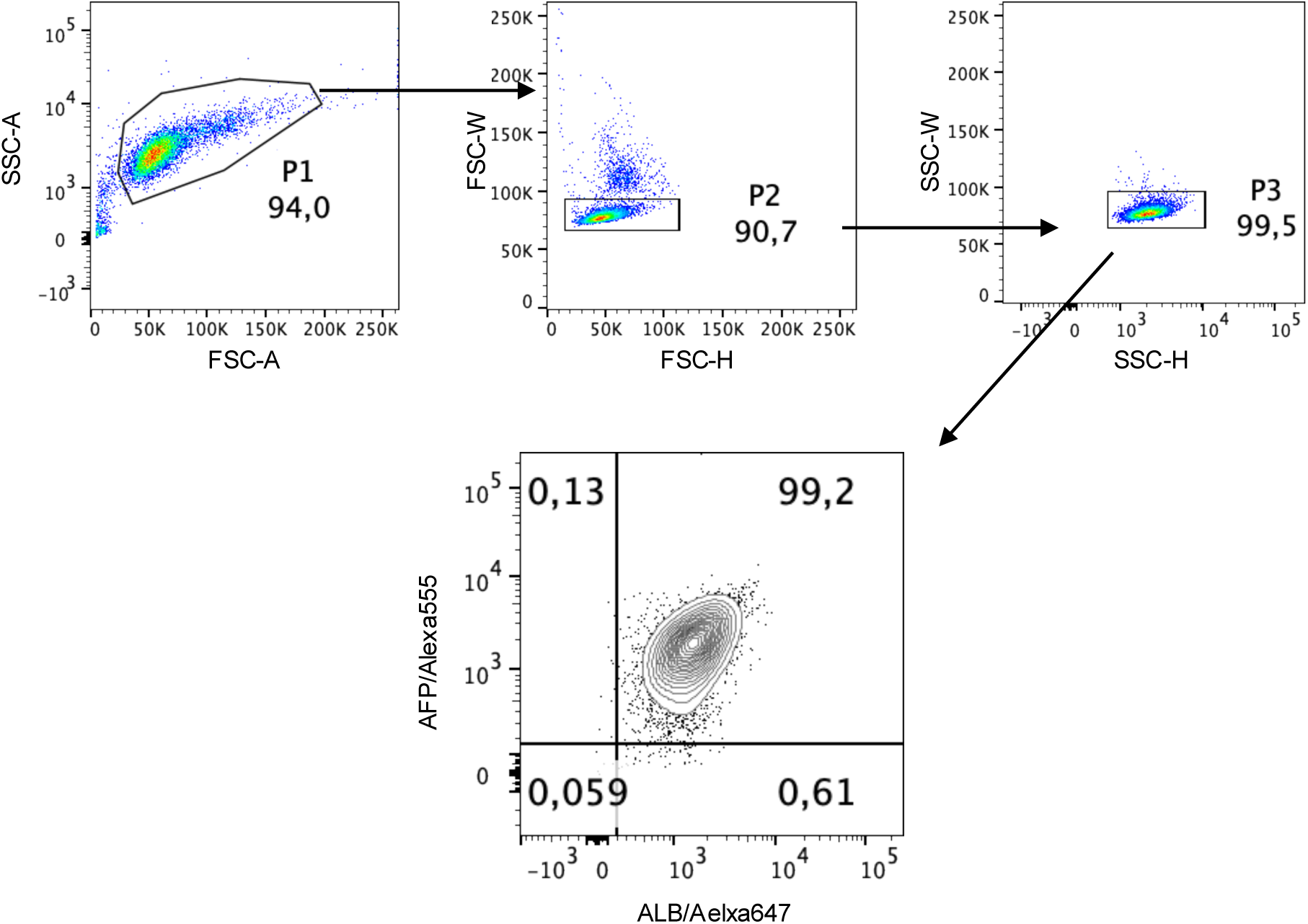
Cell gating strategy for ALB/AFP flow cytometry analysis. hPSC-derived HBs were stained with antibodies against human ALB and AFP. Gating was defined to include no more than 1% of events observed in the iso-type control.

**Supplementary Table 1.** Primary Antibodies used for Immunohistochemistry and Flow Cytometry analysis.

| Primary Antibody | Company | Product Codes | IgG Species | Dilution |
| --- | --- | --- | --- | --- |
| ALB | Bethyl | A80-129A | Goat | 1:200 |
| Ki67 | DAKO | M7240 | Mouse | 1:100 |
| AFP | DAKO | A0008 | Rabbit | 1:2000 |
| CYP3A4 | abcam | ab135813 | Rabbit | 1:200 |
| CK8/18 | abcam | ab17139 | Mouse | 1:100 |
| ASGPR | R&D | MAB4394 | Mouse | 1:100 |
| CK19 | abcam | ab52625 | Rabbit | 1:800 |
| GLUL | Sigma | HPA007316 | Rabbit | 1:200 |
| IGFBP2 | abcam | ab109284 | Rabbit | 1:100 |
| CYP2A6 | Atlas Antibodies | HPA047262 | Rabbit | 1:200 |
| KU80 | Cell Signaling Technology | 21805 | Rabbit | 1:450 |

**Supplementary Table 2.** Secondary Antibodies used for Immunohistochemistry and Flow Cytometry analysis.

| Secondary Antibody | Company | Product Codes | Dilution |
| --- | --- | --- | --- |
| IgG Donkey anti-Goat Alexa647 | Invitrogen | A21447 | 1:400 |
| IgG Donkey anti-Mouse Alexa488 | Invitrogen | A21202 | 1:400 |
| IgG Donkey anti-Rabbit Alexa555 | Invitrogen | A31572 | 1:400 |
| IgG Donkey anti-Goat Alexa488 | Invitrogen | A11055 | 1:400 |
| IgG Donkey anti-Rabbit Alexa647 | Invitrogen | A31573 | 1:400 |
| IgG Donkey anti-Mouse Alexa647 | Invitrogen | A31571 | 1:400 |

**Supplementary Table 3.**
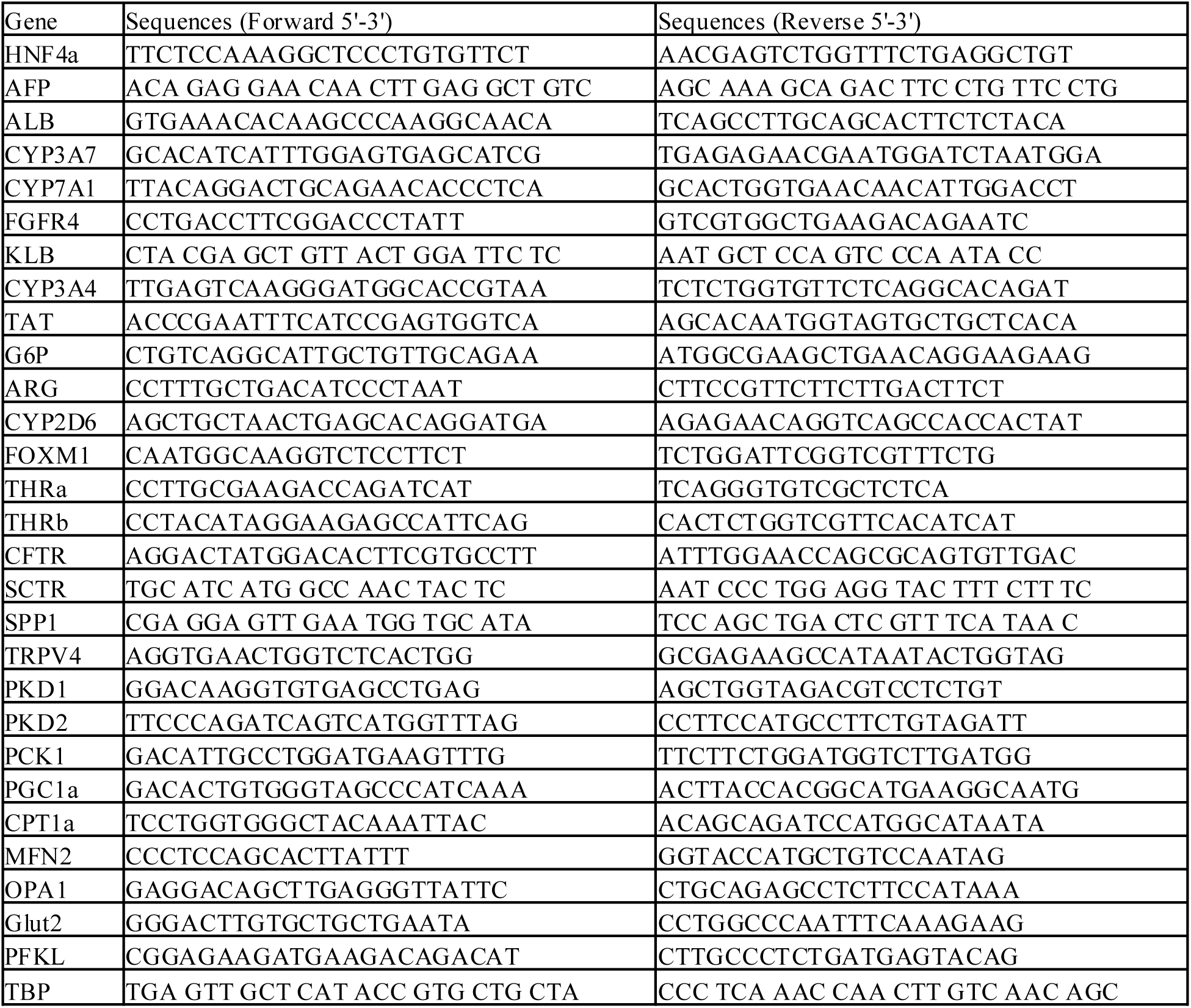
Primers used for RT-PCR analysis.

**Supplementary Table 4.** RNA controls used for RT-PCR analysis.

RNA controls used for RT-PCR analysis.
| RNA | Source | Sex | Lot | Company | Product number |
| --- | --- | --- | --- | --- | --- |
| human adult liver | normal livers pooled from 3 Asians<br>(22-64-years old) | Male | 1402003 | Clontech | 636531 |
| human fetal liver | pooled from 63 spontaneously aborted fetus,<br>aged 22-40 weeks | Male and<br>Female | 7030173 | Clontech | 636540 |
| human pancreas | normal pancreas from a 35-years old Caucasian | Male | 1703157A | Clontech | 636577 |

## Notes

### Competing Interest Statement

The authors have declared no competing interest.

## References

1 Dababneh, Y. & Mousa, O. Y. in StatPearls (2025).

2 Devarbhavi, H. et al. Global burden of liver disease: 2023 update. J Hepatol 79, 516–537 (2023). 10.1016/j.jhep.2023.03.017

3 Messina, A., Luce, E., Hussein, M. & Dubart-Kupperschmitt, A. Pluripotent-Stem-Cell-Derived Hepatic Cells: Hepatocytes and Organoids for Liver Therapy and Regeneration. Cells 9 (2020). 10.3390/cells9020420

4 Bhatia, S. N., Underhill, G. H., Zaret, K. S. & Fox, I. J. Cell and tissue engineering for liver disease. Sci Transl Med 6, 245sr242 (2014). 10.1126/scitranslmed.3005975

5 Hansel, M. C. et al. The history and use of human hepatocytes for the treatment of liver diseases: the first 100 patients. Curr Protoc Toxicol 62, 14 12 11–23 (2014). 10.1002/0471140856.tx1412s62

6 Kalra, A., Yetiskul, E., Wehrle, C. J. & Tuma, F. in StatPearls (2025).

7 Kmiec, Z. Cooperation of liver cells in health and disease. Adv Anat Embryol Cell Biol 161, III–XIII, 1–151 (2001). 10.1007/978-3-642-56553-3

8 Wilson, Z. E. et al. Inter-individual variability in levels of human microsomal protein and hepatocellularity per gram of liver. Br J Clin Pharmacol 56, 433–440 (2003). 10.1046/j.1365-2125.2003.01881.x

9 Sohlenius-Sternbeck, A. K. Determination of the hepatocellularity number for human, dog, rabbit, rat and mouse livers from protein concentration measurements. Toxicol In Vitro 20, 1582–1586 (2006). 10.1016/j.tiv.2006.06.003

10 Takahashi, K. et al. Induction of pluripotent stem cells from adult human fibroblasts by defined factors. Cell 131, 861–872 (2007). 10.1016/j.cell.2007.11.019

11 Tabar, V. et al. Phase I trial of hES cell-derived dopaminergic neurons for Parkinson’s disease. Nature 641, 978–983 (2025). 10.1038/s41586-025-08845-y

12 Sawamoto, N. et al. Phase I/II trial of iPS-cell-derived dopaminergic cells for Parkinson’s disease. Nature 641, 971–977 (2025). 10.1038/s41586-025-08700-0

13 Hogrebe, N. J., Ishahak, M. & Millman, J. R. Developments in stem cell-derived islet replacement therapy for treating type 1 diabetes. Cell Stem Cell 30, 530–548 (2023). 10.1016/j.stem.2023.04.002

14 Wang, B. et al. Functional Maturation of Induced Pluripotent Stem Cell Hepatocytes in Extracellular Matrix-A Comparative Analysis of Bioartificial Liver Microenvironments. Stem Cells Transl Med 5, 1257–1267 (2016). 10.5966/sctm.2015-0235

15 Yuan, Y., Cotton, K., Samarasekera, D. & Khetani, S. R. Engineered Platforms for Maturing Pluripotent Stem Cell-Derived Liver Cells for Disease Modeling. Cell Mol Gastroenterol Hepatol 15, 1147–1160 (2023). 10.1016/j.jcmgh.2023.01.013

16 Kajiwara, M. et al. Donor-dependent variations in hepatic differentiation from human-induced pluripotent stem cells. Proc Natl Acad Sci U S A 109, 12538–12543 (2012). 10.1073/pnas.1209979109

17 Farhan, F., Trivedi, M., Di Wu, P. & Cui, W. Extracellular matrix modulates the spatial hepatic features in hepatocyte-like cells derived from human embryonic stem cells. Stem Cell Res Ther 14, 314 (2023). 10.1186/s13287-023-03542-x

18 Huch, M. et al. In vitro expansion of single Lgr5+ liver stem cells induced by Wnt-driven regeneration. Nature 494, 247–250 (2013). 10.1038/nature11826

19 Mun, S. J. et al. Efficient and reproducible generation of human induced pluripotent stem cell-derived expandable liver organoids for disease modeling. Sci Rep 13, 22935 (2023). 10.1038/s41598-023-50250-w

20 Ma, H. et al. The nuclear receptor THRB facilitates differentiation of human PSCs into more mature hepatocytes. Cell Stem Cell 29, 1611 (2022). 10.1016/j.stem.2022.10.003

21 Reza, H. A. et al. Multi-zonal liver organoids from human pluripotent stem cells. Nature 641, 1258–1267 (2025). 10.1038/s41586-025-08850-1

22 Igarashi, R. et al. Generation of human adult hepatocyte organoids with metabolic functions. Nature 641, 1248–1257 (2025). 10.1038/s41586-025-08861-y

23 Shi, H. et al. Chemical approaches targeting the hurdles of hepatocyte transplantation: mechanisms, applications, and advances. Front Cell Dev Biol 12, 1480226 (2024). 10.3389/fcell.2024.1480226

24 Shan, J. et al. Identification of small molecules for human hepatocyte expansion and iPS differentiation. Nat Chem Biol 9, 514–520 (2013). 10.1038/nchembio.1270

25 Higashi, H. et al. Transplantation of bioengineered liver capable of extended function in a preclinical liver failure model. Am J Transplant 22, 731–744 (2022). 10.1111/ajt.16928

26 Sun, Z. et al. Hepatocyte transplantation: The progress and the challenges. Hepatol Commun 7 (2023). 10.1097/HC9.0000000000000266

27 Ogawa, M. et al. Generation of functional ciliated cholangiocytes from human pluripotent stem cells. Nat Commun 12, 6504 (2021). 10.1038/s41467-021-26764-0

28 Song, K. H., Li, T., Owsley, E., Strom, S. & Chiang, J. Y. Bile acids activate fibroblast growth factor 19 signaling in human hepatocytes to inhibit cholesterol 7alpha-hydroxylase gene expression. Hepatology 49, 297–305 (2009). 10.1002/hep.22627

29 Wang, Y. et al. An FGF15/19-TFEB regulatory loop controls hepatic cholesterol and bile acid homeostasis. Nat Commun 11, 3612 (2020). 10.1038/s41467-020-17363-6

30 Zhang, L. et al. Significance and mechanism of CYP7a1 gene regulation during the acute phase of liver regeneration. Mol Endocrinol 23, 137–145 (2009). 10.1210/me.2008-0198

31 Liu, Y. et al. Dissecting the Role of the FGF19-FGFR4 Signaling Pathway in Cancer Development and Progression. Front Cell Dev Biol 8, 95 (2020). 10.3389/fcell.2020.00095

32 Nie, Y. Z., Zheng, Y. W. & Taniguchi, H. Improving the repopulation capacity of elderly human hepatocytes by decoding aging-associated hepatocyte plasticity. Hepatology 76, 1030–1045 (2022). 10.1002/hep.32443

33 Ogawa, S. et al. Three-dimensional culture and cAMP signaling promote the maturation of human pluripotent stem cell-derived hepatocytes. Development 140, 3285–3296 (2013). 10.1242/dev.090266

34 Thorpe-Beeston, J. G., Nicolaides, K. H., Felton, C. V., Butler, J. & McGregor, A. M. Maturation of the secretion of thyroid hormone and thyroid-stimulating hormone in the fetus. N Engl J Med 324, 532–536 (1991). 10.1056/NEJM199102213240805

35 Laszlo, V. et al. Triiodothyronine accelerates differentiation of rat liver progenitor cells into hepatocytes. Histochem Cell Biol 130, 1005–1014 (2008). 10.1007/s00418-008-0482-z

36 Li, M. et al. Thyroid hormone action in postnatal heart development. Stem Cell Res 13, 582–591 (2014). 10.1016/j.scr.2014.07.001

37 Chattergoon, N. N. Thyroid hormone signaling and consequences for cardiac development. J Endocrinol 242, T145–T160 (2019). 10.1530/JOE-18-0704

38 Bogacheva, M. S., Bystriakova, M. A. & Lou, Y. R. Thyroid Hormone Effect on the Differentiation of Human Induced Pluripotent Stem Cells into Hepatocyte-Like Cells. Pharmaceuticals (Basel) 14 (2021). 10.3390/ph14060544

39 Goel, C., Monga, S. P. & Nejak-Bowen, K. Role and Regulation of Wnt/beta-Catenin in Hepatic Perivenous Zonation and Physiological Homeostasis. Am J Pathol 192, 4–17 (2022). 10.1016/j.ajpath.2021.09.007

40 Wild, S. L. et al. The Canonical Wnt Pathway as a Key Regulator in Liver Development, Differentiation and Homeostatic Renewal. Genes (Basel) 11 (2020). 10.3390/genes11101163

41 Sun, T. et al. ZNRF3 and RNF43 cooperate to safeguard metabolic liver zonation and hepatocyte proliferation. Cell Stem Cell 28, 1822–1837 e1810 (2021). 10.1016/j.stem.2021.05.013

42 Nejak-Bowen, K. & Monga, S. P. Wnt-beta-catenin in hepatobiliary homeostasis, injury, and repair. Hepatology 78, 1907–1921 (2023). 10.1097/HEP.0000000000000495

43 Plata-Gomez, A. B. et al. Hepatic nutrient and hormone signaling to mTORC1 instructs the postnatal metabolic zonation of the liver. Nat Commun 15, 1878 (2024). 10.1038/s41467-024-46032-1

44 Carson, M. D. & Nejak-Bowen, K. Wnt/beta-Catenin Signaling in Liver Pathobiology. Annu Rev Pathol 20, 59–86 (2025). 10.1146/annurev-pathmechdis-111523-023535

45 Sugimoto, A. et al. Hepatic stellate cells control liver zonation, size and functions via R-spondin 3. Nature 640, 752–761 (2025). 10.1038/s41586-025-08677-w

46 Gebhardt, R. Metabolic zonation of the liver: regulation and implications for liver function. Pharmacol Ther 53, 275–354 (1992). 10.1016/0163-7258(92)90055-5

47 Jungermann, K. & Kietzmann, T. Zonation of parenchymal and nonparenchymal metabolism in liver. Annu Rev Nutr 16, 179–203 (1996). 10.1146/annurev.nu.16.070196.001143

48 Ben-Moshe, S. & Itzkovitz, S. Spatial heterogeneity in the mammalian liver. Nat Rev Gastroenterol Hepatol 16, 395–410 (2019). 10.1038/s41575-019-0134-x

49 Halpern, K. B. et al. Single-cell spatial reconstruction reveals global division of labour in the mammalian liver. Nature 542, 352–356 (2017). 10.1038/nature21065

50 MacParland, S. A. et al. Single cell RNA sequencing of human liver reveals distinct intrahepatic macrophage populations. Nat Commun 9, 4383 (2018). 10.1038/s41467-018-06318-7

51 Aizarani, N. et al. A human liver cell atlas reveals heterogeneity and epithelial progenitors. Nature 572, 199–204 (2019). 10.1038/s41586-019-1373-2

52 Columbano, A. et al. The thyroid hormone receptor-beta agonist GC-1 induces cell proliferation in rat liver and pancreas. Endocrinology 147, 3211–3218 (2006). 10.1210/en.2005-1561

53 Sekiya, S. & Suzuki, A. Direct conversion of mouse fibroblasts to hepatocyte-like cells by defined factors. Nature 475, 390–393 (2011). 10.1038/nature10263

54 Tong, Y. F. et al. Maturity of associating liver partition and portal vein ligation for staged hepatectomy-derived liver regeneration in a rat model. World J Gastroenterol 24, 1107– 1119 (2018). 10.3748/wjg.v24.i10.1107

55 Nakano, Y. et al. Identification of a novel alpha-fetoprotein-expressing cell population induced by the Jagged1/Notch2 signal in murine fibrotic liver. Hepatol Commun 1, 215– 229 (2017). 10.1002/hep4.1026

56 Lin, C. W. et al. Hepatocyte proliferation and hepatomegaly induced by phenobarbital and 1,4-bis [2-(3,5-dichloropyridyloxy)] benzene is suppressed in hepatocyte-targeted glypican 3 transgenic mice. Hepatology 54, 620–630 (2011). 10.1002/hep.24417

57 Hasegawa, M. et al. The reconstituted ‘humanized liver’ in TK-NOG mice is mature and functional. Biochem Biophys Res Commun 405, 405–410 (2011). 10.1016/j.bbrc.2011.01.042

58 Uehara, S. et al. HepaSH cells: Experimental human hepatocytes with lesser inter-individual variation and more sustainable availability than primary human hepatocytes. Biochem Biophys Res Commun 663, 132–141 (2023). 10.1016/j.bbrc.2023.04.054

59 Kang, S. W. S. et al. A spatial map of hepatic mitochondria uncovers functional heterogeneity shaped by nutrient-sensing signaling. Nat Commun 15, 1799 (2024). 10.1038/s41467-024-45751-9

60 Yan, Y. et al. 3D bioprinting of human neural tissues with functional connectivity. Cell Stem Cell 31, 260–274 e267 (2024). 10.1016/j.stem.2023.12.009

61 Choi, S. et al. Fibre-infused gel scaffolds guide cardiomyocyte alignment in 3D-printed ventricles. Nat Mater 22, 1039–1046 (2023). 10.1038/s41563-023-01611-3

62 Di Buduo, C. A. et al. Bioprinting Soft 3D Models of Hematopoiesis using Natural Silk Fibroin-Based Bioink Efficiently Supports Platelet Differentiation. Adv Sci (Weinh) 11, e2308276 (2024). 10.1002/advs.202308276

63 Jafari, A. et al. Formulation and Evaluation of PVA/Gelatin/Carrageenan Inks for 3D Printing and Development of Tissue-Engineered Heart Valves. Advanced Functional Materials 34, 2305188 (2024). 10.1002/adfm.202305188

64 Rohani, L. et al. Stirred suspension bioreactors maintain naive pluripotency of human pluripotent stem cells. Commun Biol 3, 492 (2020). 10.1038/s42003-020-01218-3

65 Cohen, P. J. R. et al. Engineering 3D micro-compartments for highly efficient and scale-independent expansion of human pluripotent stem cells in bioreactors. Biomaterials 295, 122033 (2023). 10.1016/j.biomaterials.2023.122033

66 Dadheech, N. et al. Scale up manufacturing approach for production of human induced pluripotent stem cell-derived islets using Vertical Wheel(R) bioreactors. NPJ Regen Med 10, 24 (2025). 10.1038/s41536-025-00409-y

67 Lun, A. T., Bach, K. & Marioni, J. C. Pooling across cells to normalize single-cell RNA sequencing data with many zero counts. Genome Biol 17, 75 (2016). 10.1186/s13059-016-0947-7

68 Korsunsky, I. et al. Fast, sensitive and accurate integration of single-cell data with Harmony. Nat Methods 16, 1289–1296 (2019). 10.1038/s41592-019-0619-0

69 Subramanian, A. et al. Gene set enrichment analysis: a knowledge-based approach for interpreting genome-wide expression profiles. Proc Natl Acad Sci U S A 102, 15545– 15550 (2005). 10.1073/pnas.0506580102

70 Reimand, J. et al. Pathway enrichment analysis and visualization of omics data using g:Profiler, GSEA, Cytoscape and EnrichmentMap. Nat Protoc 14, 482–517 (2019). 10.1038/s41596-018-0103-9

